# Diversity-stability relationships become decoupled across spatial scales: a synthesis of organism and ecosystem types

**DOI:** 10.1101/2022.10.04.510879

**Authors:** Nathan I. Wisnoski, Riley Andrade, Max C.N. Castorani, Christopher P. Catano, Aldo Compagnoni, Thomas Lamy, Nina K. Lany, Luca Marazzi, Sydne Record, Annie C. Smith, Christopher M. Swan, Jonathan D. Tonkin, Nicole M. Voelker, Phoebe L. Zarnetske, Eric R. Sokol

**Author notes:** **Correspondence:** Eric R. Sokol, Nathan I. Wisnoski.

## Abstract

The relationship between biodiversity and stability, or its inverse, temporal variability, is multidimensional and complex. Temporal variability in aggregate properties, like total biomass or abundance, is typically lower in communities with higher species diversity (i.e., the diversity-stability relationship or DSR). Recent work has shown that, at broader spatial extents, regional-scale aggregate variability is also lower with higher regional diversity (in plant systems) and with lower spatial synchrony. However, it is not yet clear whether regional DSRs hold across a broad range of organisms and ecosystem types. Furthermore, focusing exclusively on aggregate properties of communities may overlook potentially destabilizing compositional shifts. To test these questions, we compiled a large collection of long-term spatial metacommunity data spanning a wide range of taxonomic groups (e.g., birds, fish, plants, invertebrates) and ecosystem types (e.g., deserts, forests, oceans). We applied a newly developed quantitative framework for jointly analyzing aggregate and compositional variability across scales. We quantified DSRs for composition and total abundance in local communities and metacommunities. At the local scale, compositional DSRs suggested that higher local (α) diversity was associated with lower variability in animal populations but higher variability in plant populations, while aggregate DSRs supported the classic stabilizing effects of diversity. Spatial synchrony differed among taxa (birds had the lowest, plants the highest), suggesting differences in stabilization by spatial processes. Spatial synchrony declined with higher diversity among sites (β) for both compositional and aggregate properties. However, at the regional (γ) scale, we found no aggregate DSR, but a positive compositional DSR. Across a broader range of taxa, our results suggest that high γ-diversity does not consistently stabilize aggregate properties at regional scales without sufficient spatial β-diversity to reduce spatial synchrony.

**Open research statement:** All data sets are accessible via the Environmental Data Initiative, and a specific data package of the data sets used in this analysis will be made publicly available (doi: pending). Citations to original sources are included in Appendix S1. Code to reproduce the analyses is found in a Zenodo archive (doi: pending) of the GitHub repository for this project (https://github.com/sokole/ltermetacommunities/tree/master/Manuscripts/MS3).

## INTRODUCTION

Temporal variability (or its inverse, stability) in the distribution and abundance of species has important implications for the maintenance of biodiversity and ecosystem functions (McCann 2000; Tilman *et al.* 2014). Ecological communities can vary through time along multiple dimensions. These dimensions include species composition (e.g., fluctuations in relative abundances) and aggregate community properties (e.g., total community abundance or biomass) that ignore composition but relate to ecosystem functioning (Micheli *et al.* 1999; Cottingham *et al.* 2001; Hillebrand *et al.* 2018; Hillebrand & Kunze 2020). Variability in aggregate community properties can be reduced through a variety of mechanisms often found in species-rich communities (Tilman 1999; McCann 2000; Ives & Carpenter 2007; Craven *et al.* 2018). This diversity-stability relationship (DSR) can arise from portfolio effects that relate to the statistical and ecological benefits of high species richness for ecosystem functioning (Doak *et al.* 1998; Tilman *et al.* 1998; Yachi & Loreau 1999; Thibaut & Connolly 2013) or from compensatory dynamics that occur when populations fluctuate asynchronously through time (Klug *et al.* 2000; Gonzalez & Loreau 2009; Brown *et al.* 2016). However, there is a pressing need to consider temporal variability at broader spatial extents to better align with the spatial scales at which ecosystem management decisions are made (e.g., ecoregions) and to incorporate the spatial dynamics of the landscape (Loreau *et al.* 2003; Wang & Loreau 2016; Wilcox *et al.* 2017; Wang *et al.* 2019; Gonzalez *et al.* 2020; Lamy *et al.* 2021).

Temporal aggregate variability at regional scales depends jointly on the variability of local communities and the degree of spatial synchrony among communities (Wang & Loreau 2014, 2016; Wang *et al.* 2019). Regional-scale fluctuations in total biomass or abundance can be reduced through a dampening of local-scale fluctuations (e.g., through local portfolio effects or compensatory dynamics) or through a reduction in spatial synchrony (Wang *et al.* 2019). Low spatial synchrony implies that community variability is weakly spatially correlated, which can dampen variability at the regional scale through a process called the spatial insurance effect (Howeth & Leibold 2010; Steiner *et al.* 2011; McGranahan *et al.* 2016; Wilcox *et al.* 2017; Catano *et al.* 2020; Wang *et al.* 2021). Spatial insurance may be strengthened by environmental heterogeneity, which generates contrasting population dynamics among communities, or by dispersal rates that are low enough to prevent spatial homogenization that could cause patches to fluctuate similarly across the metacommunity (Loreau *et al.* 2003; Gouhier *et al.* 2010; Thompson *et al.* 2015; Lamy *et al.* 2019).

Aggregate variability and species richness tend to show opposing scaling relationships with increasing spatial extent (Delsol *et al.* 2018). With increasing extent, variability declines (Wang *et al.* 2017) but richness increases (i.e., the species-area relationship) (Rosenzweig 1995). Because these changes occur simultaneously, diversity-stability relationships can be quantified across multiple spatial scales (Aragón *et al.* 2011; Wang & Loreau 2014, 2016; Wang *et al.* 2019, 2021; Gonzalez *et al.* 2020; Liang *et al.* 2022). Higher regional diversity (γ-diversity) tends to dampen variability in the aggregate properties of the whole metacommunity (Aragón *et al.* 2011; Wang & Loreau 2016; Wang *et al.* 2019), a direct scaling-up of the local aggregate DSR. This direct local-to-regional scaling has been detected in a desert grassland community, where both local and regional variability decreased as α- and γ-diversity increased, respectively (Chalcraft 2013). However, the stabilizing effect of γ-diversity across a broader survey of plant communities has received mixed support (Wilcox *et al.* 2017; Wang *et al.* 2021; Liang *et al.* 2022). Instead, growing evidence points to the regionally stabilizing effects of β-diversity, the spatial turnover in species composition, due to its direct relation to spatial synchrony (Catano *et al.* 2020; Liang *et al.* 2022; Qiao *et al.* 2022).

Regional-scale studies of the DSR have largely overlooked the relationship between diversity and compositional variability (Lamy *et al.* 2021). Quantifying compositional and aggregate variability together is important because a lack of variability in total metacommunity biomass can conceal broad-scale changes in composition (Micheli *et al.* 1999; Lamy *et al.* 2021). Furthermore, compositional studies can reveal species combinations that are important for the conservation of biomass across spatial scales (Arranz *et al.* 2022). It is not yet clear how different facets of biodiversity relate to compositional variability at local and regional scales. For example, species-rich plant communities often have higher variability in composition due to increased biotic interactions and niche partitioning (e.g., which drive compensatory dynamics), in addition to the stochastic fluctuations of communities with smaller average population sizes (Tilman 1999; Tilman *et al.* 2006; Hector *et al.* 2010; Wang *et al.* 2019). But this positive relationship between richness and compositional variability is not universal (Cottingham *et al.* 2001; Chalcraft 2013). Extending this relationship to the regional scale suggests that metacommunities with higher γ-diversity could be more compositionally variable, but empirical studies have found the lowest compositional variability at intermediate γ-diversity (Chalcraft 2013). Compositional DSRs at both local and regional spatial scales merit additional study across ecosystems and taxonomic groups to assess their generality and transferability.

Using a large compilation of long-term, spatially replicated metacommunity time series data (n=22), collected across a range of ecosystem types (e.g., deserts, forests, coral reefs) and taxonomic groups (e.g., birds, fish, plants, algae), we quantified diversity-stability relationships at multiple spatial scales. Specifically, we examined community variability (both aggregate and compositional) at both local (observational unit, such as a sampling plot) and regional (study scale, such as an LTER site) spatial scales, alongside changes in diversity, to address three questions. First (Q1), how does local diversity relate to aggregate and compositional variability within communities, and do broader relationships emerge across ecosystems and organisms? Second (Q2), does spatial heterogeneity in community composition (i.e., β-diversity) reduce spatial synchrony and stabilize both aggregate and compositional metacommunity properties at large spatial scales? And third (Q3), do aggregate and compositional variability exhibit different relationships with species diversity at the metacommunity (γ) scale? Our synthesis found wide variability among systems in compositional DSRs at the local scale, strong evidence that spatial β-diversity reduces both compositional and aggregate spatial synchrony, and the absence of an aggregate DSR, but the presence of a positive compositional DSR, at the regional scale.

## METHODS

### Data acquisition, processing, archiving

We acquired 22 data sets spanning a wide range of ecosystems and organismal groups represented by the Long-Term Ecological Research (LTER) Network, with data obtained from the Environmental Data Initiative (EDI) portal (https://portal.edirepository.org). We used time series of species assemblage data containing abundance information using the ecocomDP model (O’Brien *et al.* 2021). We kept data sets with at least five spatial and at least five temporal replicates to ensure sufficient spatial and temporal resolution. We then filtered and aggregated each data set to ensure homogeneous sampling, consistent taxonomic identification, and to make sure that each data set represented a collection of potentially interacting species (i.e., our data sets focus on competitive or “horizontal” metacommunities).

To ensure homogeneous sampling, we retained only the spatial replicates that were sampled at every time step. If some sites were not sampled annually, we retained only the portion of the data set with uninterrupted, annual temporal sampling while still meeting the criteria of a minimum of five sites and five time points. When a data set was sampled twice or more per year, we computed annual averages of species abundances to aid in comparison across data sets.

We aggregated or filtered data to ensure homogeneous taxonomic identification. For instance, our taxonomic information was provided mostly at the species level, but some data sets contained information at the genus, family, or order level. This posed an issue only when individuals were identified as belonging to a specific genus within datasets that contained several species from the same genus. For example, in certain datasets, some individuals were identified as simply belonging to a genus (e.g., “*Carex* sp.”), while most individuals were identified as a species of said genus (e.g., “*Carex phoetida*”). Such heterogeneous identification in the data suggested that species identification was inconsistent. If taxa identified at the genus level comprised a small portion of the individuals within a genus (i.e., <5% of records were identified at the genus rather than species level), we removed the genus-level data from the data set. Conversely, if >5% of individuals were identified at the genus level, we assumed this reflected inconsistent identification. In this case, we lumped taxonomic identification at the genus level. We removed all unknown species from the data sets.

We only used data sets that could represent a metacommunity by separating taxonomic groups and retaining data sets with enough shared species. For example, when a data set contained two or more distinct taxonomic groups (e.g., fish, sessile invertebrate, and algal assemblages in the Santa Barbara Coastal LTER), we treated these groups as separate data sets. Moreover, when a data set contained sites that shared <5% of species, we investigated these sites further and excluded them if they clearly did not appear to be part the same metacommunity as the majority of other sites.

To represent the salient characteristics of each data set, we plotted species accumulation curves, time series of species abundance, spatiotemporal replication, and the number of species shared between spatial replicates (Appendix S1). The data for these analyses are publicly available at EDI and listed in Table S1. All analyses were conducted with the R statistical computing environment, v. 4.1.2 (R Core Team 2022).

### Quantifying aggregate and compositional variability

To quantify variability at local and regional scales in our metacommunity data sets, we used a multiplicative partitioning framework, which decomposes regional-scale variability into the product of average local variability and spatial synchrony. Thus, higher average local variability and higher spatial synchrony both contribute to variability at the regional scale. Specifically, we partitioned metacommunity variability using aggregate (Wang & Loreau 2014, 2016) and compositional (Lamy *et al.* 2021) approaches.

For each metacommunity data set, we calculated mean local 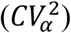 and regional 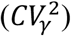 aggregate variability as well as mean local 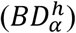 and regional 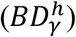 compositional variability, then partitioned variability across scales. This multiplicative partitioning yielded spatial synchrony components for both aggregate (*φ*) and compositional 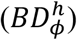 dimensions of variability in metacommunities. Values of 0 indicate no synchrony and a complete dampening of variability at regional scales, while 1 indicates no spatial stabilization. Thus, these spatial synchrony terms serve as scaling factors that link local and regional scale variability.

We compared the scaling of compositional and aggregate variability to assess the relative magnitude of spatial stabilization within and across different ecosystem types and organismal groups. We visualized the reduction in variability from local scales to the regional scale. For each metacommunity, we compared the aggregate (*φ*) versus compositional 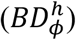 spatial synchrony among all the metacommunities, and computed Spearman’s rank correlation to quantify the association between compositional and aggregate spatial synchrony. When 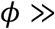 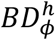, aggregate properties are weakly stabilized by space compared to compositional properties; when 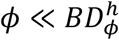, composition is weakly stabilized by space compared to aggregate properties.

### Testing multi-scale diversity-stability relationships for compositional and aggregate variability

We compared the relationship between aggregate and compositional metacommunity variability and the diversity observed at different scales in the metacommunity. We partitioned metacommunity γ-diversity with a multiplicative approach 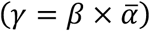, where *γ* was metacommunity richness, 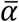 was mean local richness, and *β* was the ratio of metacommunity to local richness for each metacommunity time series in our data set. Then, 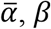, and *γ* components of diversity were averaged through time, yielding long-term estimates of local, among-site, and regional diversity for each metacommunity.

#### Question 1 (Q1): How do local-scale diversity-stability relationships differ in aggregate and compositional properties, and among ecosystems and organismal types?

To investigate local-scale relationships, we focused on within- and among-metacommunity relationships. At the local scale, we predicted that plots with higher species richness would be less variable in their aggregate properties, but that species richness would show a potentially weaker, positive relationship with compositional variability. Within a metacommunity, we computed the temporal average α-diversity of each plot and the temporal variability of total community abundance and composition. We then tested whether more species-rich plots in each metacommunity were more, or less, variable through time in their aggregate or compositional properties using linear mixed effects models in the “lme4” R package (Bates *et al.* 2015). We modeled plot-level CV or BD as the response variable and plot-level richness as the predictor. We used a random intercepts and random slopes model, such that different metacommunities can have different DSRs, while contributing to the among-group mean intercept and slope (Harrison *et al.* 2018). We computed the variance explained by the random effects using the marginal (without the random effects) and conditional (including the random effects) *R*^2^ approach for (G)LMMs (Nakagawa & Schielzeth 2013).

#### Question 2 (Q2): Does β-diversity stabilize compositional and aggregate variability at the metacommunity scale?

The scaling factors that link local variability to metacommunity variability 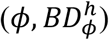 are interpreted as spatial synchrony components. To test whether these synchrony values were related to spatial heterogeneity in community composition, we compared them with a temporal average of spatial β-diversity in each metacommunity. This temporally averaged spatial β-diversity captures sustained compositional heterogeneity among plots in the metacommunity. We tested the hypothesis that metacommunities with lower average spatial β-diversity over time should have higher spatial synchrony. We predicted this relationship would be stronger for compositional than aggregate spatial synchrony.

#### Question 3 (Q3): How do aggregate and compositional diversity-stability relationships scale up to regional extents?

We then evaluated whether diversity and variability were related at the metacommunity scale. If the local scale relationships hold at regional scales, we predicted that increased γ-diversity would dampen aggregate metacommunity variability. However, the relationship with compositional metacommunity variability was predicted to be less clear given the range of patterns previously described at local and regional scales. We used ordinary least-squares linear regressions to assess the relationship between variability at the metacommunity scale and γ-diversity.

## RESULTS

Overall, we found a positive, but moderate, correlation between *φ* and 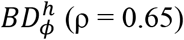, indicating that metacommunities with lower compositional spatial synchrony also tended to have less aggregate synchrony (Fig. 1). However, metacommunities differed in the relative degree of spatial synchrony in aggregate and compositional variability (i.e., deviations from the 1:1 line in Fig. 1). In other words, some metacommunities showed a greater reduction in aggregate—but not compositional—properties across scales, and vice versa. Plants and invertebrate communities tended to have the highest spatial synchrony, in both compositional and aggregate dimensions. In contrast, bird metacommunities showed consistently low spatial synchrony. Animals with slightly lower dispersal abilities (e.g., fish, zooplankton, herps) exhibited intermediate synchrony values between birds and plants.

**Fig. 1:**
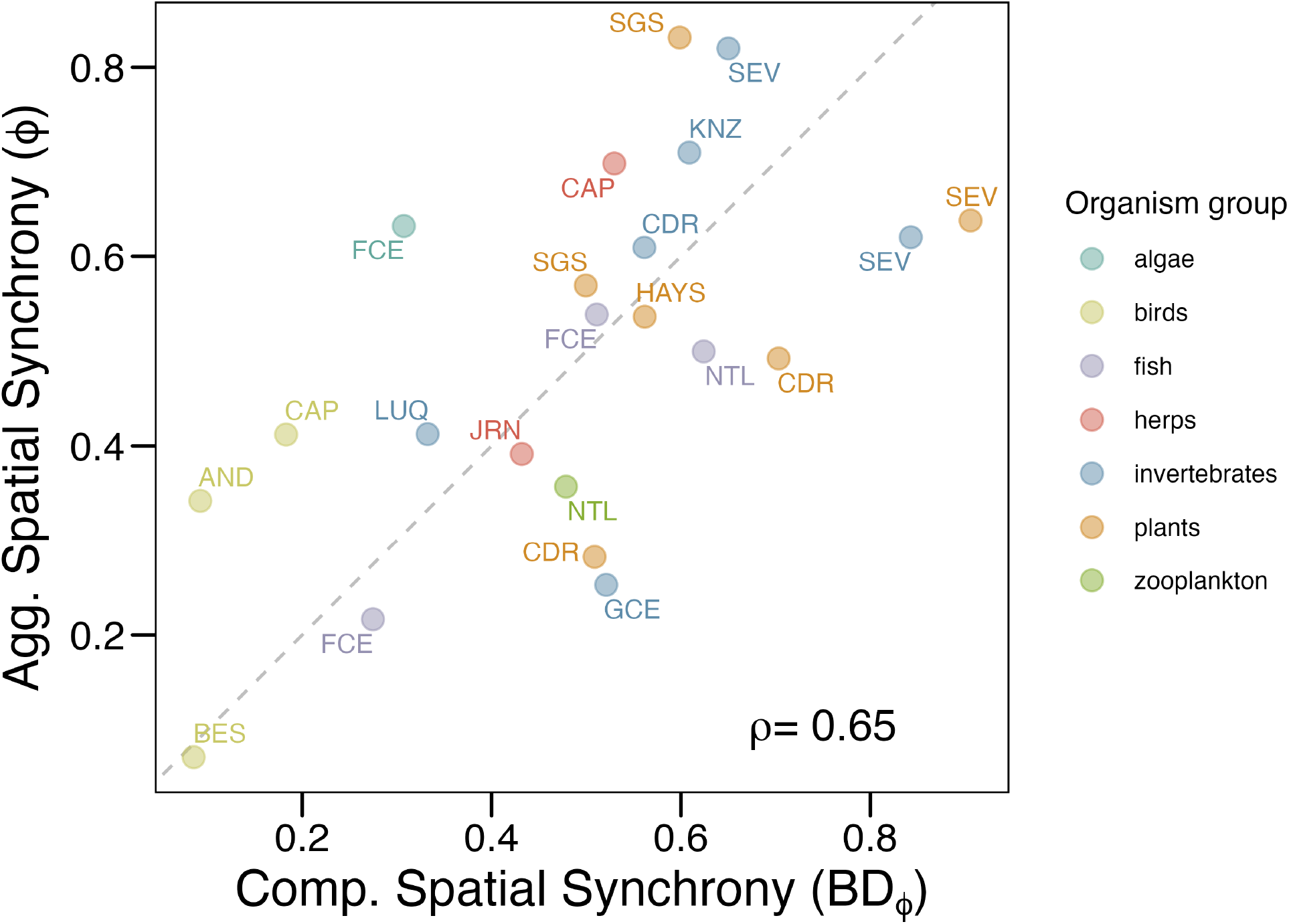
The relationship between compositional and aggregate spatial synchrony across a diverse collection of metacommunities. Compositional and aggregate synchrony were positively correlated (Spearman’s ρ =0.65) but scaled differently with space. Metacommunities above the 1:1 line had higher aggregate spatial synchrony, and thus a lower reduction in aggregate than compositional variability from local to regional scales. Metacommunities below the 1:1 line had relatively higher compositional synchrony, and thus more reduction in aggregate variability than compositional variability across scales. Colors reflect the taxonomic group of the metacommunity. AND = Andrews Forest LTER, BES = Baltimore Ecosystem Study, FCE = Florida Coastal Everglades LTER, GCE = Georgia Coastal Ecosystem LTER, CDR = Cedar Creek Ecosystem Science Reserve, HAYS = Hays Agricultural Research Center, LUQ = Luquillo Experimental Forest LTER, NTL = North Temperate Lakes LTER, JRN = Jornada Basin LTER, SGS = Shortgrass Steppe LTER, CAP = Central Arizona-Phoenix LTER, KNZ = Konza Prairie LTER, SEV = Sevilleta LTER.

We found general support for local-scale diversity-stability relationships, such that more species rich sites in the metacommunity tended to have much lower variability in total community biomass (Fig. 2A). This relationship did not hold for all communities (e.g., algae and some invertebrates), but it did emerge as a general pattern across studies as well (intercept = 0.714 ± 0.093, t = 7.642; slope = −0.019 ± 0.006, t = −2.779). However, incorporating the random effects for each dataset strongly improved the fit of the model 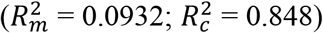. In contrast, we found a range of relationships between compositional variability and richness (Fig. 2B). Some metacommunities showed strong positive relationships between richness and compositional variability, while others showed strong negative relationships. But there was no strong effect of richness on compositional variability (intercept = 0.288 ± 0.051, t = 5.598; slope = 0.0008 ± 0.004, t = 2.07). Again, accounting for among-dataset differences drastically improved model fit 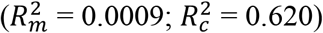, showing the importance of accounting for the variance in DSR slopes among metacommunities.

**Fig. 2:**
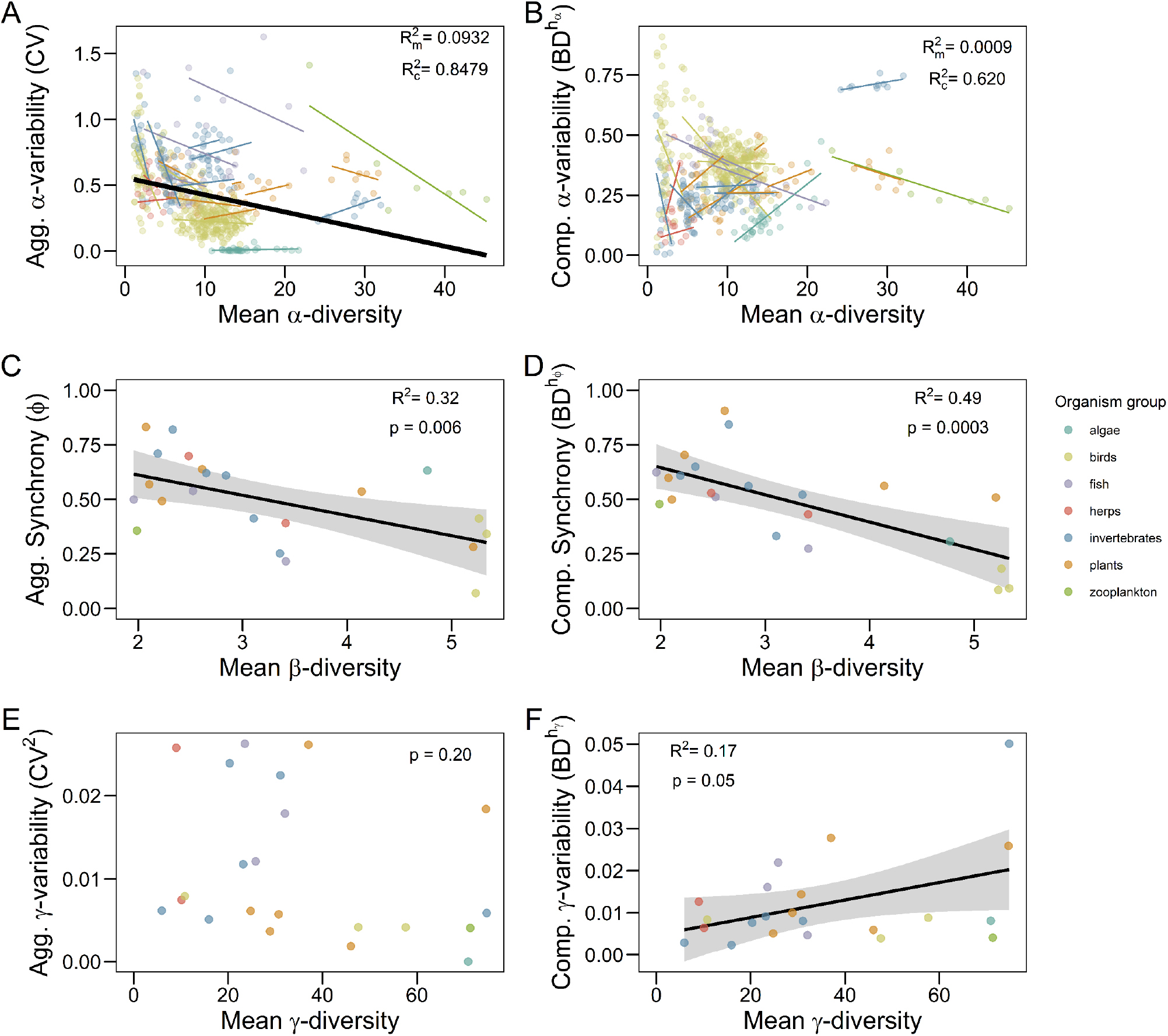
Diversity-stability relationships at the metacommunity scale. Left column: variability in aggregate properties (total abundance); right column: variability in composition (species relative abundances). A) At the local scale, aggregate variability declined with temporal mean α-diversity of each site. Bold black line represents the fixed effects of local richness on variability across studies; thinner colored lines the random slopes and intercepts accounted for by the random effects of the metacommunity dataset. The 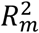 represents the variance explained by the main effects only, while the 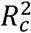 represents the variance explained by the fixed and random effects. A large difference in these values demonstrates an improvement of fit from the mixed effects model. B) Local compositional variability showed trends with α-diversity within, but not across, metacommunities. C) Mean spatial β-diversity was significantly negatively related to aggregate spatial synchrony. D) Mean spatial β-diversity was significantly negatively related to compositional spatial synchrony. E) Mean γ-diversity was not related to variability in total metacommunity abundance. F) Compositional variability at the metacommunity scale was significantly related to average γ-diversity across metacommunities. Simple linear regression fits are shown when P ≤ 0.05. Gray shading shows 95% confidence intervals. Colors are the same organismal groups as in Fig. 1.

Metacommunities with higher sustained spatial β-diversity over time exhibited both lower aggregate (Fig. 2C) and compositional (Fig. 2D) spatial synchrony. We detected a stronger relationship between β-diversity and compositional synchrony (slope = −0.125 ± 0.028 SE, F_1,20_ = 19.72, R^2^ = 0.497, p = 2.5e-4) than aggregate synchrony (slope = −0.093 ± 0.030 SE, F_1,20_ = 9.434, R^2^ = 0.321, p = 6.0e-3). Therefore, β-diversity generates spatially distinct communities that vary independently in their abundance fluctuations and in their community trajectories.

At the regional scale, we found compositional but not aggregate DSRs. Although we predicted that γ-diversity would be negatively related to aggregate variability at the metacommunity scale, we found no relationships (Fig. 2E; slope = −1.1e-4 ± 8.5e-5 SE, F_1,20_ = 1.74, R^2^ = 0.08, p = 0.2). However, γ-diversity was marginally positively related to compositional metacommunity variability (Fig. 2F; slope = 2.1e-4 ± 1e-4 SE, F_1,20_ = 4.23, R^2^ = 0.17, p = 0.05). There were not any clear patterns related to organism or ecosystem types.

## DISCUSSION

Diversity-stability relationships have been well studied at local spatial scales, but it is less clear how they scale up to regional scales through interactions between local variability and spatial synchrony. We found that local richness tended to reduce variability in aggregate properties (here, total community abundance), but we detected both positive and negative relationships between α-diversity and compositional variability. We also found large differences in spatial synchrony, with birds showing low synchrony and plants generally exhibiting high synchrony. Low spatial synchrony was associated with higher sustained spatial β-diversity, and this relationship was stronger for compositional than aggregate synchrony. For the aggregate patterns, our results suggest the stabilizing effects of α- and β-diversity may be generalizable across broader taxonomic groups, extending findings from a recent survey of multi-scale DSRs in plant communities (Liang *et al.* 2022). At larger scales, compositional metacommunity variability increased with mean γ-diversity, but we found no relationship between γ-diversity and the variability of total metacommunity abundance. Therefore, the stabilizing effects of diversity on aggregate properties shift from being driven by α-diversity at local scales to β-diversity at broader scales, but this did not translate to regionally stabilizing effects of γ-diversity. Instead, γ-diversity was associated with higher compositional variability at the regional scale, despite a stabilizing effect of β-diversity on compositional variability. Overall, our study suggests that, when analyzed across a wide range of taxonomic groups, diversity-stability relationships displayed at local scales may become decoupled at broader spatial scales.

### Taxonomic patterns in aggregate and compositional spatial synchrony

In our analysis, bird metacommunities had the lowest spatial synchrony, while plant metacommunities tended to have the highest spatial synchrony (Fig. 1). These patterns were detected in both compositional and aggregate spatial synchrony, although some metacommunities were relatively more synchronous in composition versus total community abundance. One possible explanation for this result might relate to dispersal capacity and habitat selection. Birds are the best dispersers and have the potential for habitat selection, which could promote environmental tracking (i.e., seeking favorable habitats for reproduction) across a broad-scale, heterogeneous spatial landscape (e.g., Catano *et al.* 2020). The bird metacommunity from the Baltimore Ecosystem Survey (BES) exhibited the largest reduction in variability from local to regional scales of both total abundance and composition, followed by birds in the Andrews Experimental Forest (AND) and the Central Area Phoenix (CAP) urban ecosystem (Figs. 1–2). This suggests that spatial differences in bird community dynamics, across urban and forested ecosystems, cancel each other out at the regional scale, reducing metacommunity variability. Birds in the CAP and AND ecosystems exhibited higher spatial synchrony in their total community abundances than the BES birds, suggesting a weaker spatial insurance effect than in the BES metacommunity. At the local scale, compared to other organisms, bird communities were less variable in their total abundances (Fig. 2A), but more variable in composition (Fig. 2B), which would be consistent with locally stabilizing compensation (Brown *et al.* 2016).

In contrast, plant metacommunities tended to have high spatial synchrony in both compositional and aggregate properties (Fig. 1). Such high spatial synchrony suggests that spatial processes contribute little to the reduction in variability from local to regional scales. Plants also lack habitat selection and could have a range of different passive dispersal capacities. Plants may therefore exhibit high spatial synchrony if they are not dispersal limited and communities respond similarly to environmental variability across the landscape. The spatial heterogeneity of the grassland sites included in this analysis could be insufficient to generate the high β-diversity necessary to confer regional stability. For example, several plant metacommunities have low β-diversity, which may explain why they are spatially synchronous (Fig. 2C–D). This lack of spatial insurance in plants may be due to a comparatively small spatial extent of the metacommunity compared to studies taking place at even larger spatial extents (Liang *et al.* 2022). Understanding the interplay of environmental heterogeneity and dispersal for regulating both species diversity and community dynamics at different scales could be especially relevant for conservation efforts (Andrade *et al.* 2020; Chase *et al.* 2020), and our analysis suggests there could be vast differences among taxa that must be taken into consideration before implementation.

### Diversity-stability relationships at the local scale

We observed a variety of diversity-stability relationships at the local scale. In general, we found support for the stabilizing effect of α-diversity on aggregate variability (total community biomass) (Fig. 2A). However, there was variation in DSR slopes among organismal groups. For example, several invertebrate communities did not follow a negative relationship between diversity and aggregate variability, but instead showed a positive relationship. Fish, zooplankton, bird, and most plant communities supported a stabilizing effect of biodiversity. Relative to the numerous studies analyzing DSRs in plant communities, our broader analysis suggests there could be deviations from predicted DSRs that depend on the organisms in the community.

We found heterogeneous relationships between diversity and compositional variability (Fig. 2B). Overall, there was no significant relationship between richness and compositional variability across all metacommunities, but there were strong relationships among sites within metacommunities. For example, many (but not all) invertebrate communities showed strong negative relationships such that communities with higher α-diversity were less variable in their composition over time, as we observed with bird and fish communities. In contrast, algae and plant communities tended to show positive relationships between mean α-diversity and compositional variability (Fig. 2B). This positive relationship is consistent with the predictions established in other plant communities (Tilman 1999; Tilman *et al.* 2006; Hector *et al.* 2010; Wang *et al.* 2019). Our broader synthesis suggests that this positive relationship may not translate to communities of other organisms, especially animal communities.

### β-diversity reduces spatial synchrony

The strongest and most consistent relationship that emerged in our study was the negative relationship between mean spatial β-diversity and spatial synchrony (Fig. 2C–D). This result shows that this prediction holds across metacommunities spanning a range of ecosystem types and organismal groups, including species with different dispersal capabilities. Moreover, because spatial synchrony in both composition and total abundance declined with β-diversity, this suggests aggregate variability declined across scales due to spatially distinct community dynamics, rather than non-compositional factors like asynchronous fluctuations in community size (Lamy *et al.* 2021). This result also supports the prediction that β-diversity is important for reducing aggregate γ-variability (Wang & Loreau 2014), due to its effects on reducing compositional synchrony (Wang & Loreau 2016). Recent empirical work in temperate forests across northern China further supports this prediction by showing that reducing spatial synchrony is more important than reducing local variability for stabilizing aggregate properties at broad spatial scales (Qiao *et al.* 2022). In applied contexts where the management aim is to stabilize aggregate properties at broad scales (e.g., reducing fluctuations in the overall productivity of a multi-species fishery), then the long-term maintenance of spatial ecological processes that promote asynchrony in the metacommunity could be desirable (Harrison *et al.* 2020). More generally, the maintenance of β-diversity may have a range of benefits that are relevant for multidimensional (e.g., diversity, productivity, biomass, functioning) conservation aims at large spatial scales (Socolar *et al.* 2016).

### Diversity-stability relationships at the regional scale

Our results suggest a transition in how DSRs scale up from local to metacommunity scales. While α-diversity tends to reduce aggregate fluctuations at the local scale, we found no evidence that higher γ-diversity reduced the variability of total metacommunity abundance (Fig. 2E). Although γ-diversity reduced aggregate variability at the regional scale in recent syntheses of plant biodiversity experiments focusing on within-ecosystem patterns (Wang *et al.* 2019, 2021; Liang *et al.* 2022), our results suggest that this relationship may not hold more broadly across a wider collection of organisms and ecosystems. The absence of a regional DSR across ecosystems may occur because different numbers of species may be needed to stabilize different ecosystems, making γ-diversity a poor predictor of variability. For example, some metacommunities with low γ-diversity had low aggregate variability (Fig. 2E), possibly indicating that a few well adapted species may be sufficient to stabilize regional abundances, while other high γ-diversity systems may include species that are vulnerable to disturbances or environmental extremes that reduce regional abundances. Other dimensions of community structure, such as evenness (Craven *et al.* 2018; Valencia *et al.* 2020) or the spatial synchrony of richness (Walter *et al.* 2021), may also be relevant for community variability beyond species richness alone. While we analyzed total community abundances, rather than total biomass, diversity can stabilize a range of aggregate properties of communities (Lehman & Tilman 2000; Gonzalez & Loreau 2009; Loreau *et al.* 2021), and our results are consistent with other syntheses that found weak relationships between diversity and biomass production at metacommunity scales (Wilcox *et al.* 2017). Thus, in contrast to theoretical predictions that γ-diversity should stabilize aggregate properties at broad spatial scales, empirical systems may exhibit other sources of variability that dilute the strength of this predicted relationship among ecosystems.

Although there were important system-specific nuances, we found a positive relationship between γ-diversity and compositional metacommunity variability (Fig. 2F). This observation is consistent with decades of earlier work predicting that complex systems with greater diversity may tend toward instability at the species level, captured here by our compositional variability metric (May 1973; Tilman 1999; Ives & Carpenter 2007). Interestingly, while aggregate properties shifted from a consistent negative relationship between α-diversity and local variability to a lack of relationship at the regional scale, compositional variability shifted from a heterogeneous collection of local scale DSRs towards a more consistent pattern at the regional scale. Thus, our analysis shows that spatial processes at the metacommunity scale do not obscure this relationship, but rather make it stronger across metacommunities relative to the heterogeneity of local-scale compositional diversity-stability relationships (Fig. 2B).

### Joint scaling of diversity and variability across space

It is important to keep in mind that both diversity and variability change in tandem with spatial scale. Despite the stabilizing effects on aggregate variability of α-diversity on local scales and β-diversity on spatial synchrony, γ-diversity was not stabilizing even though α- and β-diversity increase γ-diversity. Likewise, for compositional variability, the effects of α-diversity were mixed and, even though β-diversity reduced spatial synchrony, higher γ-diversity was still associated with higher variability. These differences in the ability of local versus spatial components to maintain diversity or reduce variability suggest that it is crucial to consider the specific facets of diversity and variability in a metacommunity to understand the scaling of diversity-stability relationships more broadly.

## ACKNOWLEDGEMENTS

We thank the many researchers who collected the data used in this publication and the research sites and funding agencies that support long-term research. We acknowledge support from LTER Metacommunities Synthesis Group (NSF DEB#1545288), administered by the Long-Term Ecological Research Network Office (LNO) located at the National Center for Ecological Analysis and Synthesis (NCEAS), University of California, Santa Barbara. Research was also facilitated by the National Ecological Observatory Network (NEON), sponsored by the National Science Foundation and administered by Battelle Memorial Institute. Logistical and data management support was provided by the Environmental Data Initiative (EDI). JDT is supported by a Rutherford Discovery Fellowship administered by the Royal Society Te Apārangi (RDF-18-UOC-007) and Bioprotection Aotearoa and Te Pūnaha Matatini, both Centres of Research Excellence funded by the Tertiary Education Commission, New Zealand. ERS was partially supported on NSF DEB grant 1655593.

## Author Contributions

NIW led the analysis, made the figures, and wrote the first draft of the manuscript. All authors contributed to data processing, conceptualization of the study, analysis, figures, and writing/edits.

The authors declare no conflicts of interest.

## REFERENCES

Andrade, R., Franklin, J., Larson, K.L., Swan, C.M., Lerman, S.B., Bateman, H.L., et al. (2020). Predicting the assembly of novel communities in urban ecosystems. Landsc. Ecol.

Aragón, R., Oesterheld, M., Irisarri, G. & Texeira, M. (2011). Stability of ecosystem functioning and diversity of grasslands at the landscape scale. Landsc. Ecol., 26, 1011–1022.

Arranz, I., Fournier, B., Lester, N.P., Shuter, B.J. & Peres-Neto, P.R. (2022). Species compositions mediate biomass conservation: The case of lake fish communities. Ecology, 103, e3608.

Bates, D., Mächler, M., Bolker, B. & Walker, S. (2015). Fitting linear mixed-effects models using lme4. J. Stat. Softw., 67, 1–48.

Brown, B.L., Downing, A.L. & Leibold, M.A. (2016). Compensatory dynamics stabilize aggregate community properties in response to multiple types of perturbations. Ecology, 97, 2021–2033.

Catano, C.P., Fristoe, T.S., LaManna, J.A. & Myers, J.A. (2020). Local species diversity, β-diversity and climate influence the regional stability of bird biomass across North America. Proc. R. Soc. B Biol. Sci., 287, 20192520.

Chalcraft, D.R. (2013). Changes in ecological stability across realistic biodiversity gradients depend on spatial scale. Glob. Ecol. Biogeogr., 22, 19–28.

Chase, J.M., Jeliazkov, A., Ladouceur, E. & Viana, D.S. (2020). Biodiversity conservation through the lens of metacommunity ecology. Ann. N. Y. Acad. Sci., 1469, 86–104.

Cottingham, K.L., Brown, B.L. & Lennon, J.T. (2001). Biodiversity may regulate the temporal variability of ecological systems. Ecol. Lett., 4, 72–85.

Craven, D., Eisenhauer, N., Pearse, W.D., Hautier, Y., Isbell, F., Roscher, C., et al. (2018). Multiple facets of biodiversity drive the diversity–stability relationship. Nat. Ecol. Evol., 2, 1579–1587.

Delsol, R., Loreau, M. & Haegeman, B. (2018). The relationship between the spatial scaling of biodiversity and ecosystem stability. Glob. Ecol. Biogeogr., 27, 439–449.

Doak, D.F., Bigger, D., Harding, E.K., Marvier, M.A., O’Malley, R.E. & Thomson, D. (1998). The statistical inevitability of stability-diversity relationships in community ecology. Am. Nat., 151, 264–276.

Gonzalez, A., Germain, R.M., Srivastava, D.S., Filotas, E., Dee, L.E., Gravel, D., et al. (2020). Scaling-up biodiversity-ecosystem functioning research. Ecol. Lett., 23, 757–776.

Gonzalez, A. & Loreau, M. (2009). The causes and consequences of compensatory dynamics in ecological communities. Annu. Rev. Ecol. Evol. Syst., 40, 393–414.

Gouhier, T.C., Guichard, F. & Gonzalez, A. (2010). Synchrony and stability of food webs in metacommunities. Am. Nat., 175, E16–E34.

Harrison, H.B., Bode, M., Williamson, D.H., Berumen, M.L. & Jones, G.P. (2020). A connectivity portfolio effect stabilizes marine reserve performance. Proc. Natl. Acad. Sci., 117, 25595–25600.

Harrison, X.A., Donaldson, L., Correa-Cano, M.E., Evans, J., Fisher, D.N., Goodwin, C.E.D., et al. (2018). A brief introduction to mixed effects modelling and multi-model inference in ecology. PeerJ, 6, e4794.

Hector, A., Hautier, Y., Saner, P., Wacker, L., Bagchi, R., Joshi, J., et al. (2010). General stabilizing effects of plant diversity on grassland productivity through population asynchrony and overyielding. Ecology, 91, 2213–2220.

Hillebrand, H. & Kunze, C. (2020). Meta-analysis on pulse disturbances reveals differences in functional and compositional recovery across ecosystems. Ecol. Lett., 23, 575–585.

Hillebrand, H., Langenheder, S., Lebret, K., Lindström, E., Östman, Ö. & Striebel, M. (2018). Decomposing multiple dimensions of stability in global change experiments. Ecol. Lett., 21, 21–30.

Howeth, J.G. & Leibold, M.A. (2010). Species dispersal rates alter diversity and ecosystem stability in pond metacommunities. Ecology, 91, 2727–2741.

Ives, A.R. & Carpenter, S.R. (2007). Stability and diversity of ecosystems. Science, 317, 58–62.

Klug, J.L., Fischer, J.M., Ives, A.R. & Dennis, B. (2000). Compensatory dynamics in planktonic community responses to pH perturbations. Ecology, 81, 387–398.

Lamy, T., Wang, S., Renard, D., Lafferty, K.D., Reed, D.C. & Miller, R.J. (2019). Species insurance trumps spatial insurance in stabilizing biomass of a marine macroalgal metacommunity. Ecology, 100, e02719.

Lamy, T., Wisnoski, N.I., Andrade, R., Castorani, M.C.N., Compagnoni, A., Lany, N., et al. (2021). The dual nature of metacommunity variability. Oikos, 130, 2078–2092.

Lehman, C.L. & Tilman, D. (2000). Biodiversity, stability, and productivity in competitive communities. Am. Nat., 156, 534–552.

Liang, M., Baiser, B., Hallett, L.M., Hautier, Y., Jiang, L., Loreau, M., et al. (2022). Consistent stabilizing effects of plant diversity across spatial scales and climatic gradients. Nat. Ecol. Evol., 1–7.

Loreau, M., Barbier, M., Filotas, E., Gravel, D., Isbell, F., Miller, S.J., et al. (2021). Biodiversity as insurance: from concept to measurement and application. Biol. Rev., 96, 2333–2354.

Loreau, M., Mouquet, N. & Gonzalez, A. (2003). Biodiversity as spatial insurance in heterogeneous landscapes. Proc. Natl. Acad. Sci., 100, 12765–12770.

May, R.M. (1973). Stability and complexity in model ecosystems. Princeton University Press, Princeton, NJ.

McCann, K.S. (2000). The diversity-stability debate. Nature, 405, 228–233.

McGranahan, D.A., Hovick, T.J., Elmore, R.D., Engle, D.M., Fuhlendorf, S.D., Winter, S.L., et al. (2016). Temporal variability in aboveground plant biomass decreases as spatial variability increases. Ecology, 97, 555–560.

Micheli, F., Cottingham, K.L., Bascompte, J., Bjørnstad, O.N., Eckert, G.L., Fischer, J.M., et al. (1999). The dual nature of community variability. Oikos, 85, 161–169.

Nakagawa, S. & Schielzeth, H. (2013). A general and simple method for obtaining R2 from generalized linear mixed-effects models. Methods Ecol. Evol., 4, 133–142.

O’Brien, M., Smith, C.A., Sokol, E.R., Gries, C., Lany, N., Record, S., et al. (2021). ecocomDP: A flexible data design pattern for ecological community survey data. Ecol. Inform., 101374.

Qiao, X., Geng, Y., Zhang, C., Han, Z., Zhang, Z., Zhao, X., et al. (2022). Spatial asynchrony matters more than alpha stability in stabilizing ecosystem productivity in a large temperate forest region. Glob. Ecol. Biogeogr., 31, 1133–1146.

R Core Team. (2022). R: A language and environment for statistical computing. R Foundation for Statistical Computing, Vienna, Austria. URL https://www.R-project.org/.

Rosenzweig, M.L. (1995). Species diversity in space and time. Cambridge University Press, Cambridge, UK.

Socolar, J.B., Gilroy, J.J., Kunin, W.E. & Edwards, D.P. (2016). How should beta-diversity inform biodiversity conservation? Trends Ecol. Evol., 31, 67–80.

Steiner, C.F., Stockwell, R.D., Kalaimani, V. & Aqel, Z. (2011). Dispersal promotes compensatory dynamics and stability in forced metacommunities. Am. Nat., 178, 159–170.

Thibaut, L.M. & Connolly, S.R. (2013). Understanding diversity-stability relationships: Towards a unified model of portfolio effects. Ecol. Lett., 16, 140–150.

Thompson, P.L., Beisner, B.E. & Gonzalez, A. (2015). Warming induces synchrony and destabilizes experimental pond zooplankton metacommunities. Oikos, 124, 1171–1180.

Tilman, D. (1999). The ecological consequences of changes in biodiversity: a search for general principles. Ecology, 80, 1455–1474.

Tilman, D., Isbell, F. & Cowles, J.M. (2014). Biodiversity and ecosystem functioning. Annu. Rev. Ecol. Evol. Syst., 45, 471–493.

Tilman, D., Lehman, C.L. & Bristow, C.E. (1998). Diversity-stability relationships: statistical inevitability or ecological consequence? Am. Nat., 151, 277–282.

Tilman, D., Reich, P.B. & Knops, J.M.H. (2006). Biodiversity and ecosystem stability in a decade-long grassland experiment. Nature, 441, 629–632.

Valencia, E., de Bello, F., Galland, T., Adler, P.B., Lepš, J., E-Vojtkó, A., et al. (2020). Synchrony matters more than species richness in plant community stability at a global scale. Proc. Natl. Acad. Sci., 201920405.

Walter, J.A., Shoemaker, L.G., Lany, N.K., Castorani, M.C.N., Fey, S.B., Dudney, J.C., et al. (2021). The spatial synchrony of species richness and its relationship to ecosystem stability. Ecology, 102, e03486.

Wang, S., Lamy, T., Hallett, L.M. & Loreau, M. (2019). Stability and synchrony across ecological hierarchies in heterogeneous metacommunities: linking theory to data. Ecography, 42, 1200–1211.

Wang, S. & Loreau, M. (2014). Ecosystem stability in space: α, β and γ variability. Ecol. Lett., 17, 891–901.

Wang, S. & Loreau, M. (2016). Biodiversity and ecosystem stability across scales in metacommunities. Ecol. Lett., 19, 510–518.

Wang, S., Loreau, M., Arnoldi, J.-F., Fang, J., Rahman, K.A., Tao, S., et al. (2017). An invariability-area relationship sheds new light on the spatial scaling of ecological stability. Nat. Commun., 8, 15211.

Wang, S., Loreau, M., Mazancourt, C. de, Isbell, F., Beierkuhnlein, C., Connolly, J., et al. (2021). Biotic homogenization destabilizes ecosystem functioning by decreasing spatial asynchrony. Ecology, 102, e03332.

Wilcox, K.R., Tredennick, A.T., Koerner, S.E., Grman, E., Hallett, L.M., Avolio, M.L., et al. (2017). Asynchrony among local communities stabilises ecosystem function of metacommunities. Ecol. Lett., 20, 1534–1545.

Yachi, S. & Loreau, M. (1999). Biodiversity and ecosystem productivity in a fluctuating environment: the insurance hypothesis. Proc. Natl. Acad. Sci. U. S. A., 96, 1463–1468.

