## Appendix S1 for "Diversity-stability relationships become decoupled across spatial scales: a synthesis of organism and ecosystem types"

October 4, 2022

#### 1 Data overview

This document briefly describes the data cleaning, subsetting, and aggregation methods specific to each dataset. We plot species accumulation curves, spatio-temporal variation in the number of taxa observed, the spatio-temporal sampling effort, and the number of taxa shared by each pairwise combination of plots within the study. For each dataset, we evaluated whether all sites were observed with equal effort in all years of study and we document any years, sites, sampling occasions, or species that were dropped from the dataset prior to analysis. We also document the method used to aggregate the data to an annual value for each species at each site. We visually evaluated the species accumulation curves and temporal patters in species abundances to screen for changes in sampling methods or data recording methods over the course of the study that could affect results. For example, a sudden jump in the number of taxa in a given year could indicate that taxa were recorded to a finer resolution (not that more taxa were actually present at the site). We also checked to be sure an abundance of zero was recorded for each absence (i.e. we made sure zeros were filled in for site-years in which a species was not observed). We removed species codes that obviously represent 'unknown' species.

#### 2 Marine datasets

##### 2.1 gce-mollusc

Data were downloaded from several EDI packages (approx. one per year of study). Data are in Figure 1.

### 3 Freshwater datasets

#### 3.1 fce-diatoms

Data were obtained from the PI. Metadata and citation can be found on EDI (Gaiser, 2017). Diatom abundance collected from 800m x 800m Principal Sampling Units (hereafter called ?sites?) distributed across the Florida Everglades. Data retained for analysis include 192 total diatom taxa from 30 sites sampled at least once per year for 7 consecutive years. Full data set included 367 diatom taxa and 171 sites; however, not all sites were sampled every year. We retained sites from Shark River Slough (SRS) and Taylor Slough (TSL) of Everglades National Park that were sampled in 7 consecutive years. Diatom abundance was aggregated as yearly mean of four samples collected per year (three samples for 2006). All unknown species and species that were not detected over the time series at these sites were removed. Data are shown in Figure 2

#### 3.2 fce-fish

Data were obtained from the PI, and are cataloged on EDI (Rehage, 2017). Catch per unit effort (CPUE) was calculated as (count/distance)*100 For each species, the CPUE was aggregated by summing the CPUE measured in each bout (i.e., replicate) within each Creek Number — River — YEAR — Season combination. The format was re-arranged to reflect the aggregated CPUE ('total CPUE') per season as columns and mean CPUE column was created by averaging the CPUE across the three seasons. NOTE: After the first preliminary examination of the aggregated CPUE values the mean CPUE was ignored due to the unbalanced observations across the years and seasons. So, only the observations in the Wet and Dry season were considered. Dry season: In this set, only the sites within RB were considered to create a balanced design for the analysis. Thus, this set encompassed a longer temporal range with a cost of less spatial replication. NOTE: Also, YEAR 2004-2005 and 2011 were eliminated due to incomplete representation across the sites Wet season: In this set both RB and TB were considered but only after 2010. Thus, this set has a wider spatial replication but a shorter temporal range. NOTE: The difference in the spatial replication between seasons is related to the limitation of the sampling approach (electrofishing) which is limited to low salinity conditions. Salinity threshold for electrofishing is often reached in TB during the Dry season (i.e., TB is considered the estuarine portion of the Shark River System). Data from the dry season are shown in Figure 3. Data from the wet season are shown in Figure 4.

##### 3.3 ntl-zooplankton

The data were downloaded from the EDI Data Portal (Center for Limnology, 1983). Samples were taken via vertical tows at fortnightly intervals on a minimum of five occasions per year (range = 5 - 18 occasions per year). Density was recorded as number of individuals per liter for each taxa, integrated volumetrically over the water column. Many taxa are identified to species level, but some are identified to genus level. Lake 'Tr' was only sampled in one year and was assumed to be the same as lake 'TR'. The initial year (1981) was removed from analysis because only five of the seven lakes were sampled. We additionally removed 165 records with missing or unknown taxa designations. Data were aggregated annually for each taxa in each lake by taking the maximum density observed in a tow sampling occasion. Data are shown in Figure 5.

##### 3.4 ntl-fish

**species codes from the southern lakes were not dropped, so the metadata indicates there are 81 total species even though there are under 60.** Data on fish abundance were downloaded from the EDI Data Portal (Magnuson et al., 2010). Two types of gill nets (VGN and VGN127) and the method "ESHOCK" were rarely used, and thus removed from the dataset.The gear method "ESHOCK" was also rarely used and a follow up call is needed to determine how best to handle those data. They are currently included. Catch per unit effort (CPUE) was calculated for each species in each lake per year across all gear types (electrofishing, gill nets, baited traps) as the total catch divided by the total effort. We should double check with dataset contacts to be sure we did this correctly. 88 rows containing unidentified species were removed. Also, the two bog lakes (CB, TB) had 1-3 species each and were removed. Only data from the northern lakes were included. Data are shown in Figure 6.

### 4 Terrestrial datasets

#### 4.1 cdr-plants

Data were downloaded from EDI (Tilman, 2018). Annual censuses for dataset are incomplete after 2004, so we only consider data until 2004. We used data from control plots only, from all four sites (A, B, C, and D). Fields A, B, and C were considered together as a single dataset, while D was considered a separate datasets because of its unique fire history. We cleaned taxonomic data by cleaning clear mistakes (such as ‘carex sp.’ instead of ‘Carex sp.’), removed non-taxonomic entities (“Miscellaneous litter”) and non-plant taxa (‘Fungi’, and ‘Mosses & lichens’). We also lumped or dropped certain taxonomic information. In particular, we lumped taxonomic information to the genus level when more than 1% (more than a proportion of 0.01) of the biomass of a genus was not identified to the species level. For example, *Cyperus* species are usually identified to the genus level (“Cyperus sp.”) rather than species level (“*Cyperus schweinitzii*”), with a minority of the biomass (less than 10%) identified at the specific level. We therefore consider only genus information for *Cyperus*. We dropped taxonomic information when within a certain genus, biomass is identified to the genus level. For example, in the genus *Viola*, more than 99.8% of biomass was determined to the species level. We therefore dropped from the dataset the instances when biomass is assigned to ’Viola sp.’. Data are shown in figures 7 and 8.

#### 4.2 hays-plants

Data were downloaded from Ecological Archives (Adler et al., 2007). We removed unknown species, species categorized as ‘short grass’ and ‘Fragment’, and ‘Bare ground’. Moreover, we identified seven issues with species identification. First, we removed individuals identified a the genus level for *Ambrosia*, *Oxalis*, and *Solidago*. In the genera more than 95% of individuals were identified at the species level. Second, we ‘lumped’ to genus level, species belonging to the general *Allium*, *Chamaesyce*, *Opuntia*, *Polygala*. In these genera, more than 5% of counts were identified at the genus level. Finally, we only retained the 14 plots with continuous replication between 1938 and 1973. Data are shown in Figure 9.

##### 4.3 sev-plants

Data were downloaded from EDI (Muldavin, 2015). This dataset has a consistent spatio-temporal sampling: quadrats, within four transects (N,S,R,V) within two plots, all at one site, sampled twice a year (spring and fall). There is, however, a third plots (plot number 3) that contains four new transect (A,B,C,D), which contain 5 - rather than 10 - quadrats each. Moreover, the replicates have been occasionally censused in winter, but not consistently through the years. Therefore, we removed plot 3 and winter censuses. We then removed NAs in species counts and species identity, and summed species counts across the 10 quadrats contained in each transect. Finally, the "DATE" variable is defined as "year.month" (hence, 2000.5 refers to year 2000 in May). We chose to have season 2 (spring) correspond to May, and season 3 (Fall) correspond to September. Data are shown in Figure 10.

##### 4.4 sgs-plants

Data were downloaded from EDI (Stapp, 2013). Plant community composition on the three grassland and three shrubland small mammal trapping webs (hereafter called 'sites'; n = 6). Vegetation measurements were made once per year, usually in mid-July. Percent canopy cover of each plant species was estimated visually in 30 0.10-m2 Daubenmire quadrats on each web. We aggregated plant species percent cover data at the ?site? scale because each year, transects and plot locations were determined based on a randomization procedure (3 trap stations were chosen randomly within each site from a list of 12 permanent points, where transects with random orientations were centered on each trap station location). Trapping web ?31W? was removed because it was sampled in only 2 years (2006 and 2007). Samples for year 2007 were removed because not all sites were sampled in 2007. Species codes that did not identify plant species were removed (litter, bare ground, etc.). Species codes that were otherwise inconsistent (different cases, naming conventions, etc.) were reconciled so that all species were identified by unique 4 letter codes. Observations for codes that did not represent plants, or plants were unknown or not resolved to species were removed prior to analysis. Data are shown in Figure 11.

##### 4.5 sev-arthropods

The data were downloaded from the EDI Data Portal Data are shown in Figure 12.

##### 4.6 sev-grasshoppers

The data were downloaded from the EDI Data Portal (Lightfoot, 2010). We retained data from only two habitats (Black grama and Creosotebush) that shared many species, and presented 20 years of temporal replication. This data were collected twice a year, and spatial replication included site, transect, and web. Population (count) data is structured, being collected across sex, age, and substrate. We summed numbers across webs, sex, age, and substrate. Data are shown in Figure 13.

##### 4.7 cdr-grasshoppers

Data were accessed on EDI (Knops, 2018). We removed sites '28' and '11', which were added to the sampling after 1989. Then, we summed the number of individuals across all life stages and all months. We summed across months even if in 2003, June and August samples were lost for some fields. However, as the documentation reports, "The total counts for these fields were augmented by proportional additions from remaining samples by John Haarstad and crew". Taxonomic information is available for only for the *Acrididae* family before 1994. We lumped taxonomic identifications to genus level for *Conocephalus*, *Scudderia*, and it Tetrix, as too high a proportion of individuals from these genera were not identified at the species level. Similarly, some individuals belonging to the *Melanoplus* genus were identified at the genus level only. We removed these records, as they comprised only 6% of the entire counts in the *Melanoplus* genus. Data are shown in Figure 14.

##### 4.8 knz-grasshoppers

Data on grasshopper abundance (1996-2015) were downloaded from EDI (Joern, 2018). We retained 13 spatial replicates which provide continuous temporal replicates from 1995 to 2013 (2012 is missing). We lumped taxonomic information level whenever the counts at the genus level made up more than 5% of the total individuals counted. We did not do this lumping for *Melanopus* spp.

because it is a hyperdiverse genus. In this case, we dropped all *Melanopus* records identified at the genus level. Finally, we average count data across a year, because some spatial replicates could occasionally contain more observations within a year. Data are shown in Figure 15.

##### 4.9 luq-snails

Data were downloaded from EDI (Willig, 2010). We averaged number of snails across runs ("Run.ID" in dataset) and seasons ("Seasons" in dataset). There were no codes for unknown or non-living taxa. Data are shown in Figure 16.

##### 4.10 jrn-lizards

The data were downloaded from the EDI Data Portal (Whitford et al., 1991). This was a mark-recapture study. Pitfall traps were opened for two weeks four times per year (quarterly). The monthly samples from 1990 and 1991 were removed. Individual lizards were identified and the number of unique individuals per site per year were summed. Two sites that were established five years after the start of the study (SUMM and NORT) were excluded. We don't have a key to the species codes. Data are shown in Figure 17. The cumulative number of taxa was still increasing at the end of the time series, although with only 20 species total one or two species introductions could cause this pattern.

##### 4.11 cap-herps

The data were downloaded from the EDI Data Portal (Bateman and Childers, 2018). Data are shown in Figure 18. This is a visual encounter survey where herpetofauna observations were nested by 3 plots within 3 transects per site. Each site represents the reach level (each reach level site is composed of 3 transects with equal area sampling efforts). Surveys from 2012 and March were dropped to standardize sampling efforts temporally. Unidentified species were dropped, species code is the scientific name of species. Taxon count was calculated as the maximum abundance per year in any one of the sampling events (with 3 sampling events per reach in April/May, June/July, and September/October). The final community dataset prepared for analysis has 7 sites representative of the reach level metacommunity dynamics (3 transects sampled per reach) from 2013-2017 with a total of 18 observed species across all sites.

##### 4.12 and-birds

The data were downloaded from the EDI Data Portal (Hadley, 2017). We used data from the first five years of study (2009 - 2013) because six counts per season were conducted in those years. We summed the counts of new individuals observed within the closest distance radius (¿ 50 m) of the points during the 10-minute interval for on each sampling occasion, and used the maximum number of individuals of each species recorded at each point as the abundance value for that year. Data are shown in Figure 19.

##### 4.13 cap-birds

Data were obtained from EDI (Bateman et al., 2017). Data are shown in Figure 20. This is a point count study where birds were observed (seen or heard) for 15 minutes within a 40 m fixed radius. Each site represents one point count. Only ESCA point counts are included. Data from 2017 were dropped because not all sites were sampled. Four point count sites (M-9, V-18, X-8, and V-16) were dropped due to uneven sampling across years. Unidentified species accounted for less then 2% of the total data and were dropped. Taxon count was calculated as the maximum abundance per year during the spring month's sampling events (with 3 sampling events per site between March, April, and May).

##### 4.14 bes-birds

Data were downloaded from EDI (Nilon and Brodsky, 2017). We dropped all observations in the distance catagory "FT" and all observations not identified to species. We only used sites with surveys every year from 2005-2009. When there was multiple surveys per year for a single site, the abundance counts were aggregated to the maximum observed count from any survey for each species at the respective site. We only included plots in which at least one bird was observed in each year of study. Data are shown in (Figure 21).

#### References


Adler, P. B., Tyburczy, W. R., and Lauenroth, W. K. (2007). Long-term mapped quadrats from kansas prairie: demographic information for herbaceaous plants. *Ecology*, 88(10):2673.

Bateman, H. and Childers, D. (2018). Long-term monitoring of herpetofauna along the Salt and Gila Rivers in and near the greater Phoenix metropolitan area, ongoing since 2012. Environmental Data Initiative. http://dx.doi.org/10.6073/pasta/2999764ec0527e78d44cc9ea9fc14bf0. Dataset accessed 5/15/2018.

Bateman, H., Childers, D., Katti, M., Shochat, E., and Warren, P. (2017). Point-count bird censusing: long-term monitoring of bird abundance and diversity in central arizona-phoenix, ongoing since 2000. Environmental Data Initiative. http://dx.doi.org/10.6073/pasta/201add557165740926aab6e056db6988. Dataset accessed 5/15/2018.

Center for Limnology, N. L. (1983). North Temperate Lakes LTER: Zooplankton - Trout Lake Area 1982 - current. Environmental Data Intiative. http://dx.doi.org/10.6073/pasta/c866e3663bae76388f63233a5fdfb3d4.

Gaiser, E. (2017). Relative abundance diatom data from periphyton samples collected for the Comprehensive Everglades Restoration Plan (CERP) study (FCE) from February 2005 to November 2014. Environmental Data Initiative. https://doi.org/10.6073/pasta/cb0f7e88d28075a6ff1f59d008bb732c. Dataset obtained from PI 1/30/2017.

Hadley, S. J. K. F. (2017). Forest-wide bird survey at 183 sample sites the Andrews Experimental Forest from 2009-present. Environmental Data Initiative. http://dx.doi.org/10.6073/pasta/b75f34d8de6fcfa59c882bb1acf96226. Dataset accessed 4/25/2018.

Joern, A. (2018). CGR02 sweep sampling of grasshoppers on Konza Prairie LTER watersheds (1982-present). Environmental Data Initiative. https://doi.org/10.6073/pasta/aec67f5d71d14cd39fe8b6b34b4719f4. Dataset accessed 11/2/2018.

Knops, J. (2018). Core old field grasshopper sampling:successional dynamics on a resampled chronosequence. Environmental Data Initiative. https://doi.org/10.6073/pasta/239b3023d75d83e795a15b36fac702e2. Dataset accessed 11/2/2018.

Lightfoot, D. (2010). Long-term Core Site Grasshopper Dynamics for the Sevilleta National Wildlife Refuge, New Mexico (1992-2013). Environmental Data Intiative. https://doi.org/10.6073/pasta/c1d40e9d0ec610bb74d02741e9d22576.

Magnuson, J., Carpenter, S., and Stanley, E. (2010). North Temperate Lakes LTER: Fish Abundance 1981 - current. Environmental Data Intiative. http://dx.doi.org/10.6073/pasta/7ed3313d08fbfc92656262b977508340.

Muldavin, E. (2015). Pinon-Juniper (Core Site) Quadrat Data for the net Primary Production Study at the Sevilleta National Wildlife Refuge, New Mexico (2003-present ). Environmental Data Intiative. https://doi.org/10.6073/pasta/f19a93a5c6879a9380c394dabbf2db3a.

Nilon, C. and Brodsky, C. (2017). Biodiversity - fauna - bird survey. Environmental Data Initiative. https://doi.org/10.6073/pasta/c790bd0b1bdde870b5b1e7f631d3d38e. Dataset accessed 7/11/2018.

Rehage, J. (2017). Seasonal electrofishing data from rookery branch and tarpon bay, everglades national park (fce) from november 2004 to present. Environmental Data Initiative. http://dx.doi.org/10.6073/pasta/ed3febe89ff59f68ae2aedf6c87b7eff. Dataset obtained from PI 5/15/2018.

Stapp, P. (2013). SGS-LTER Long-Term Monitoring Project: Vegetation Cover on Small Mammal Trapping Webs on the Central Plains Experimental Range, Nunn, Colorado, USA 1999 -2006, ARS Study Number 118. Environmental Data Intiative. https://doi.org/10.6073/pasta/f6ecfd9d99c9cc83e47469d10b620cfb. Accessed Nov 7, 2018.

Tilman, D. (2018). Plant aboveground biomass data: Long-Term Nitrogen Deposition: Population, Community, and Ecosystem Consequences. Environmental Data Initiative. https://doi.org/10.6073/pasta/93b27926879861815ebb021bfd6f14ae. Dataset accessed 5/16/2018.

Whitford, W., Lightfoot, D., and Anderson, J. (1991). Lizard pitfall trap data (LTER-II, LTER-III). Environmental Data Intiative.http://dx.doi.org/10.6073/pasta/411d2828c578c4777218fce541cc4291.

Willig, M. R. (2010). El Verde Grid long-term invertebrate data. Environmental Data Intiative. https://doi.org/10.6073/pasta/ec88f3dd4ed8e172802b52ff3bb82aa8.

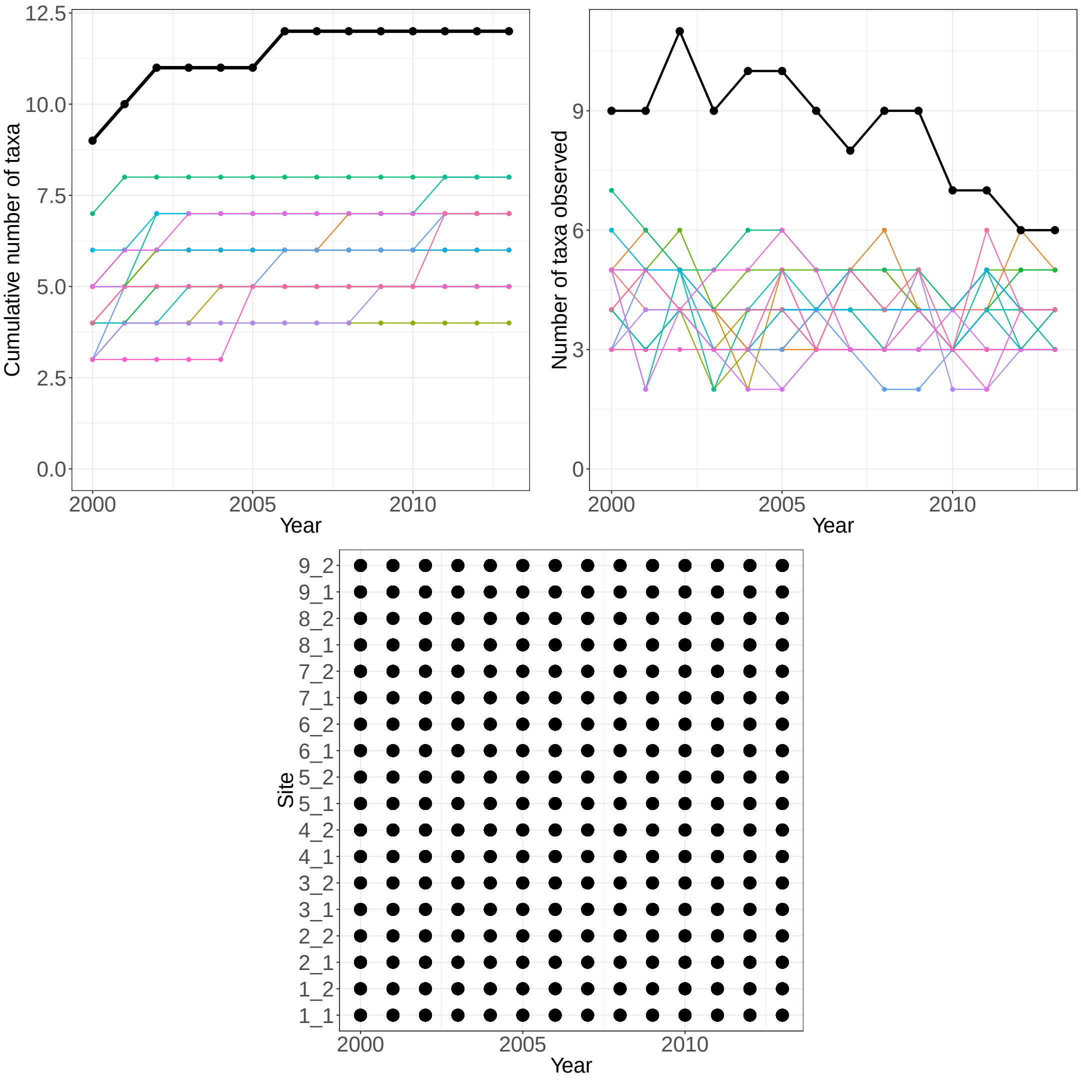


Figure 1: **GCE-mollusc:** Species accumulation curves (top left), annual richness (top right), and sampling effort (bottom) for 11 mollusc taxa observed at Georgia Coastal Ecosystems LTER. The black lines represent total site-level values across all plots.

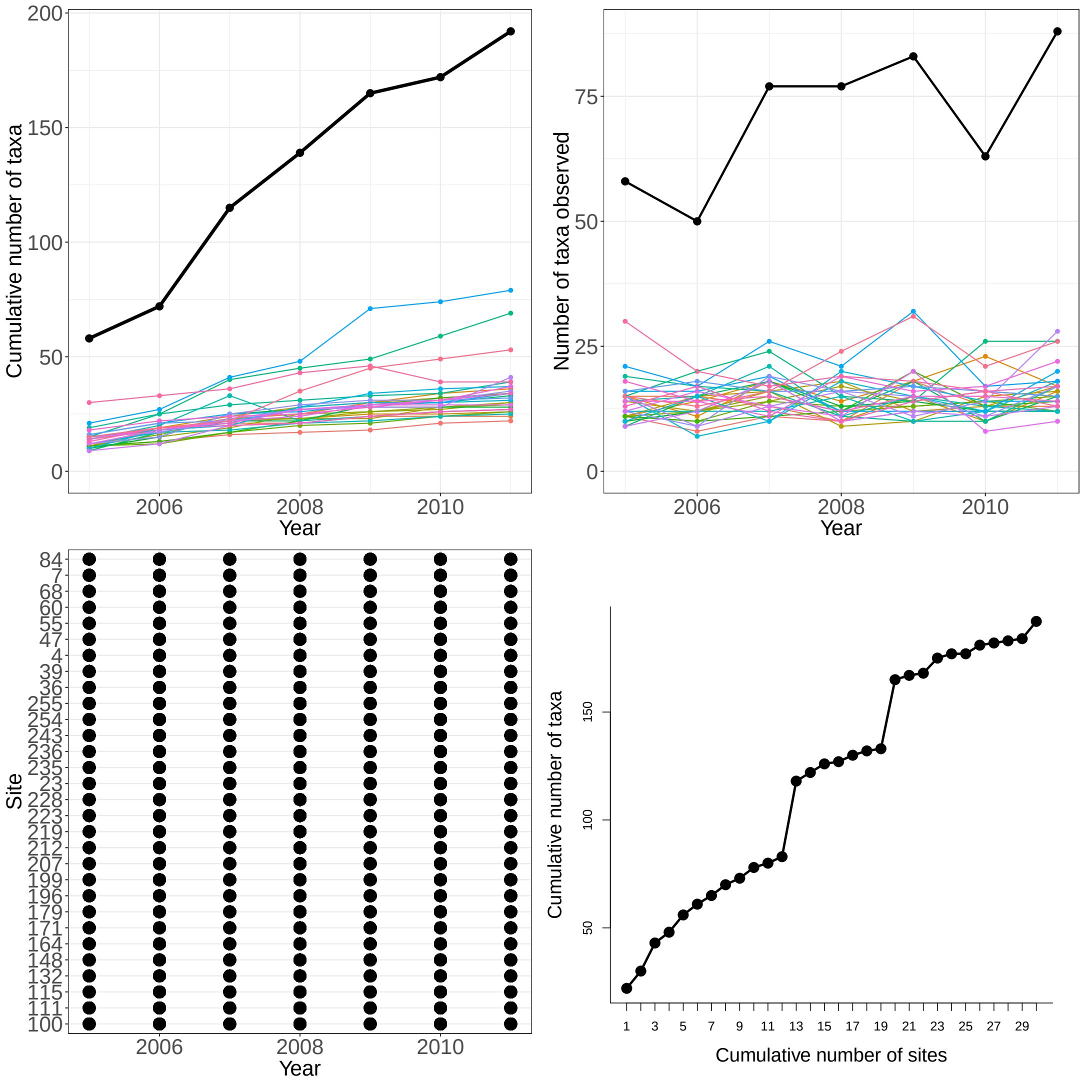


Figure 2: **FCE-diatoms:** Species accumulation curves (top left), annual richness (top right), and sampling effort (bottom) for diatom taxa observed at Florida Coastal Everglades LTER. The black lines represent total site-level values across all plots.

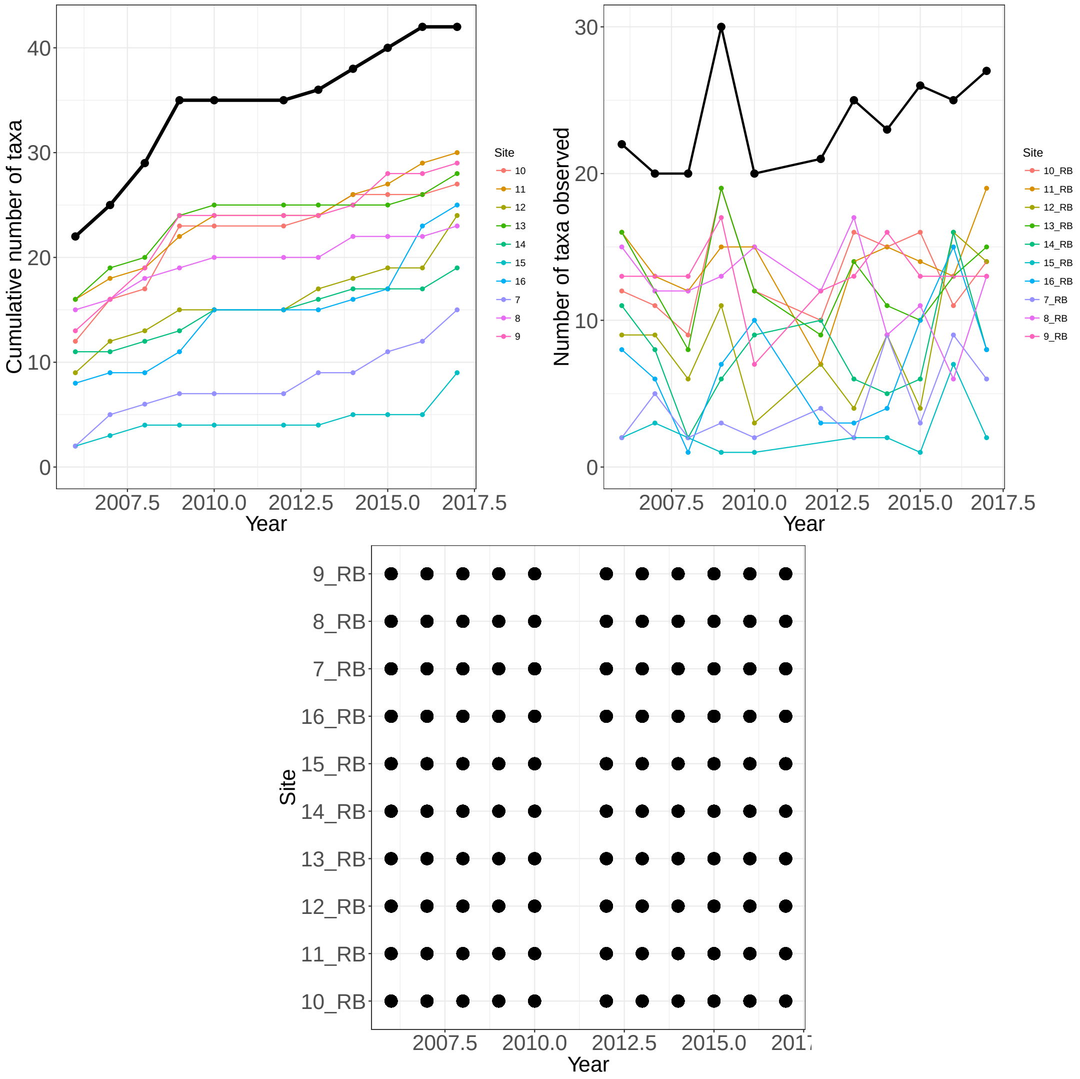


Figure 3: **FCE-fish dry season:** Species accumulation curves (top left), annual richness (top right), and sampling effort (bottom) for fish taxa observed at the Florida Coastal Everglades . The black lines represent total site-level values across all plots.

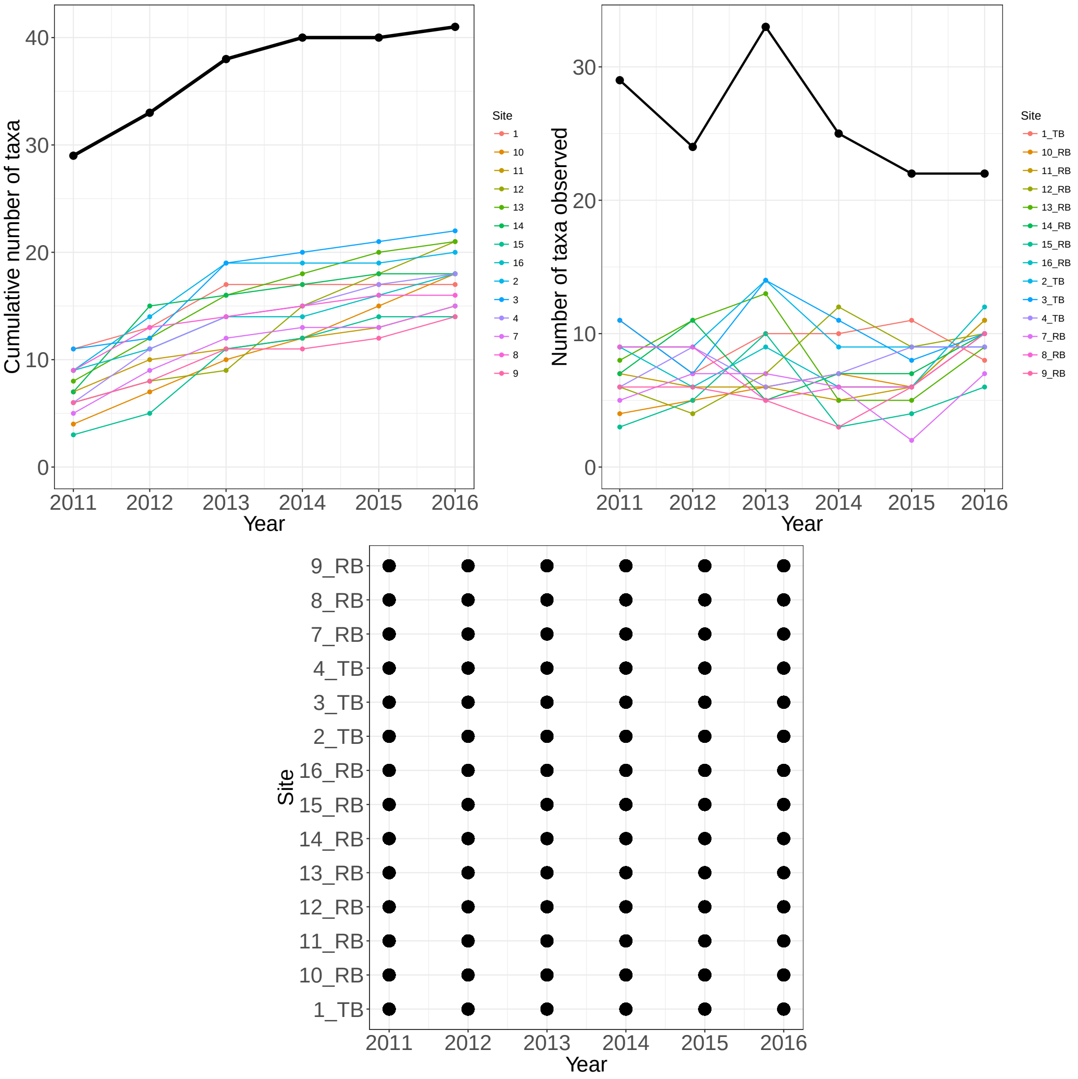


Figure 4: **FCE-fish wet season:** Species accumulation curves (top left), annual richness (top right), and sampling effort (bottom) for fish taxa observed at the Florida Coastal Everglades . The black lines represent total site-level values across all plots.

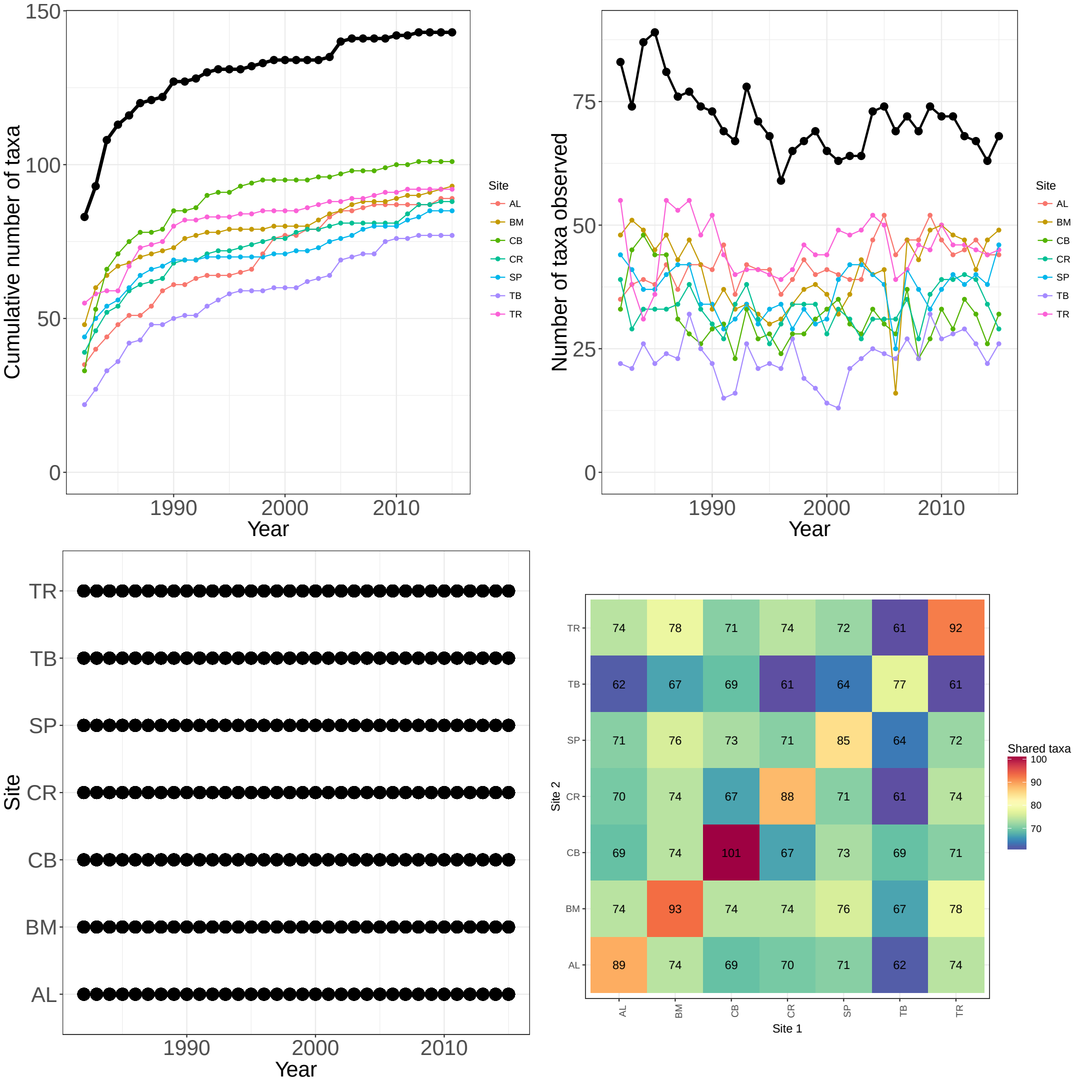


Figure 5: **NTL-zooplankton:** Species accumulation curves (top left), annual richness (top right), and sampling effort (bottom) for 143 zooplankton taxa observed at 7 sites North Temperate Lakes LTER . The black lines represent total site-level values across all plots. The plot of the species accumulation curve failed because one site was sampled in only one year, and will be fixed once that site is removed.

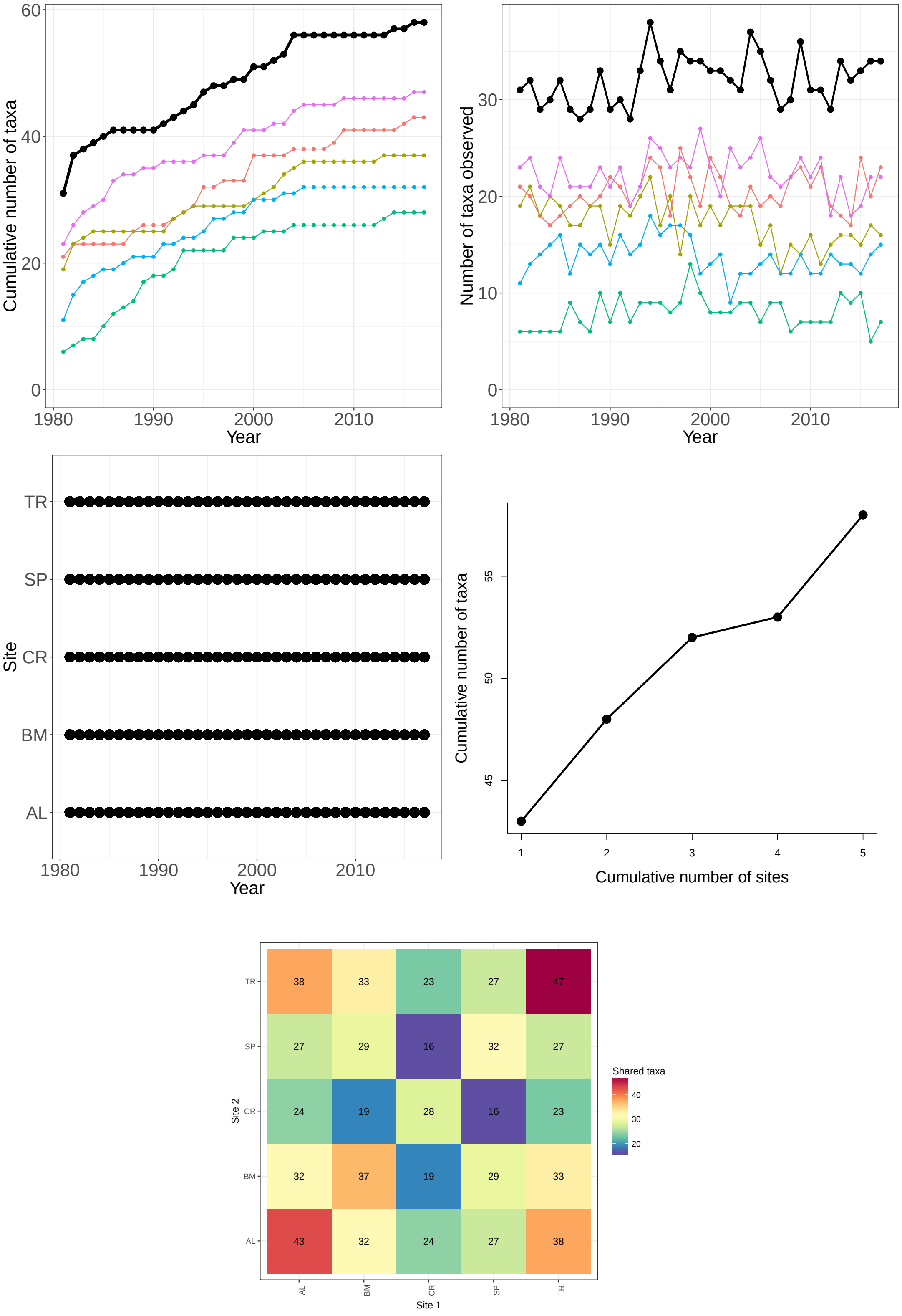


Figure 6: **NTL-fish:** Species accumulation curves (top left), annual richness (top right), and sampling effort (bottom) for 81 fish species observed at 9 lakes in the North Temperate Lakes LTER (1995-2016). The black lines represent total site-level values across all plots.

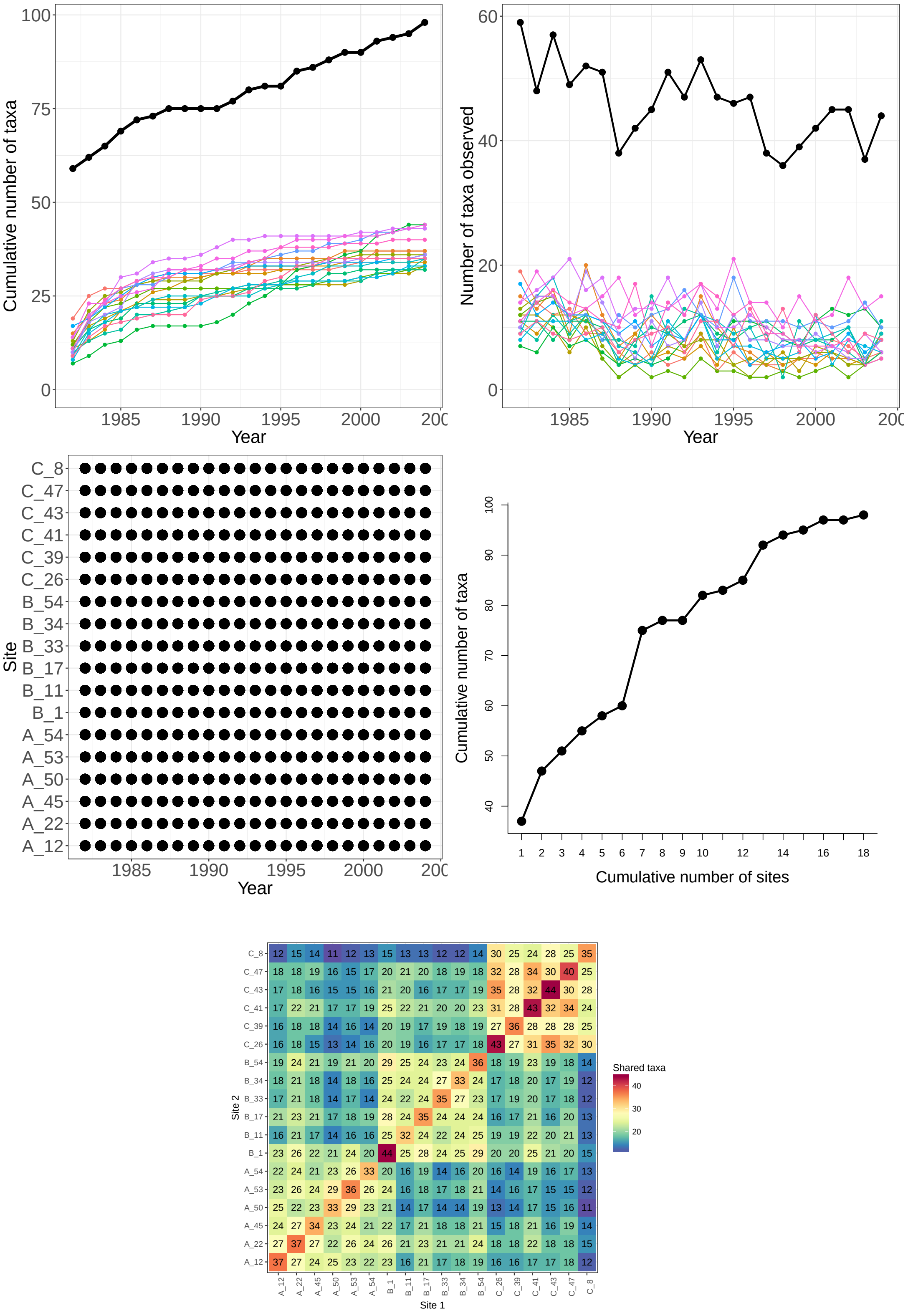


Figure 7: **CDR-plants, A, B, and C fields:** Species accumulation curves (top left), annual richness (top right), and sampling effort (bottom) for plant species observed in the A, B, and C fields at Cedar Creek. The black lines represent total site-level values across all plots.

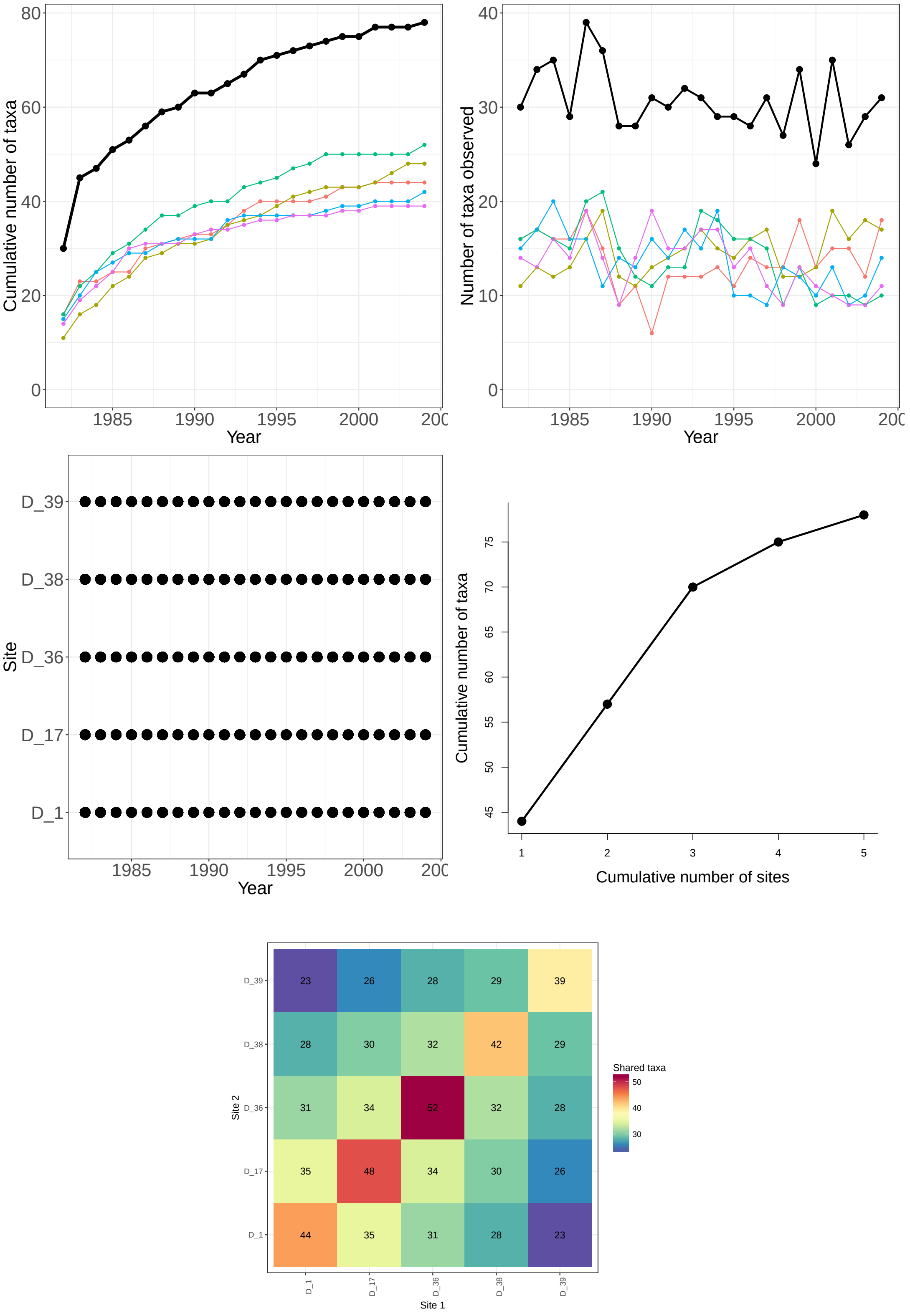


Figure 8: **CDR-plants, D field:** Species accumulation curves (top left), annual richness (top right), and sampling effort (bottom) for plant species observed in the D field at Cedar Creek. The black lines represent total site-level values across all plots.

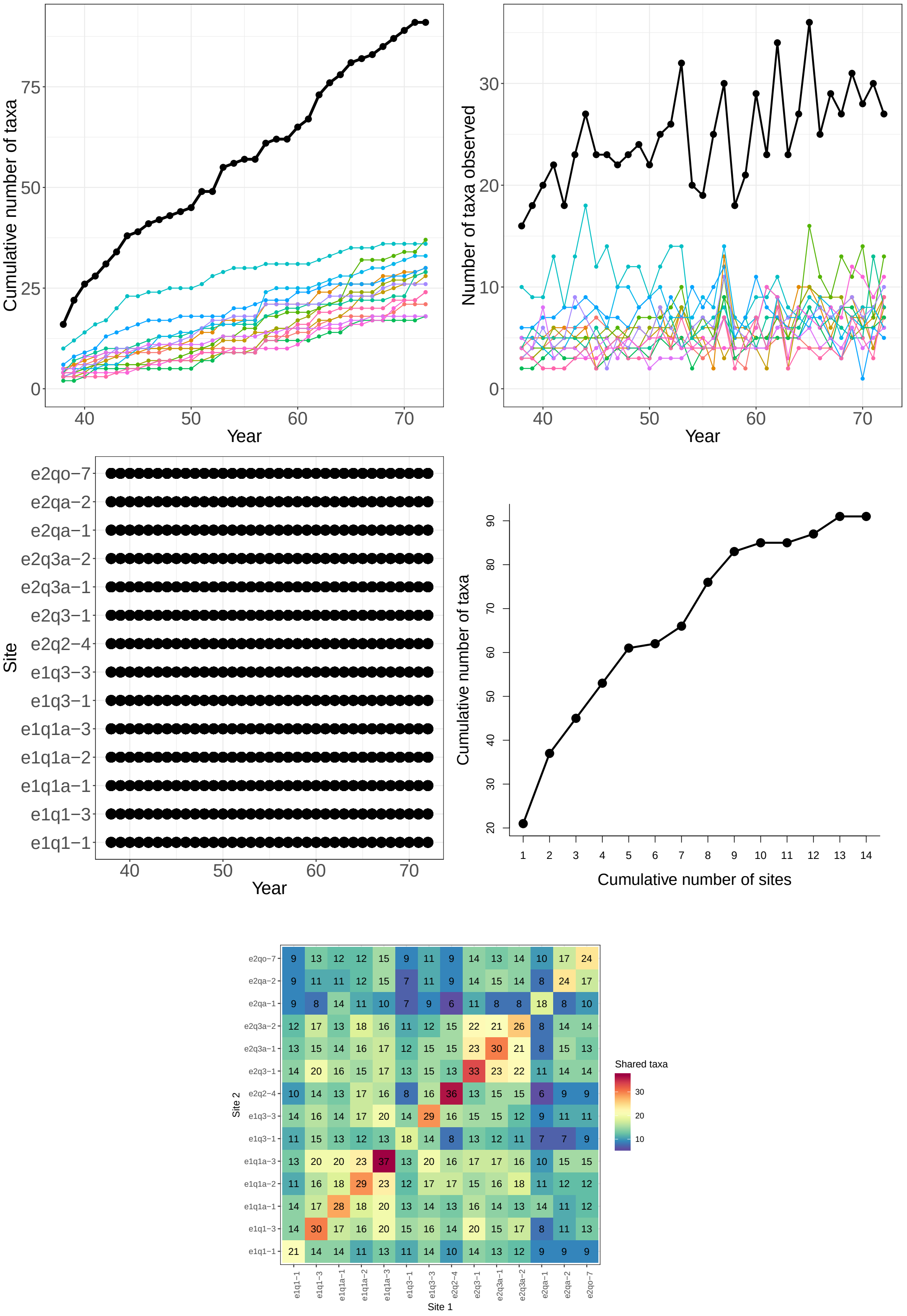


Figure 9: **hays-plants:** Species accumulation curves (top left), annual richness (top right), and sampling effort (bottom) for mixed-grass prarie plant species observed at long-term monitoring plots in Hays, Kansas. The black lines represent total site-level values across all plots.

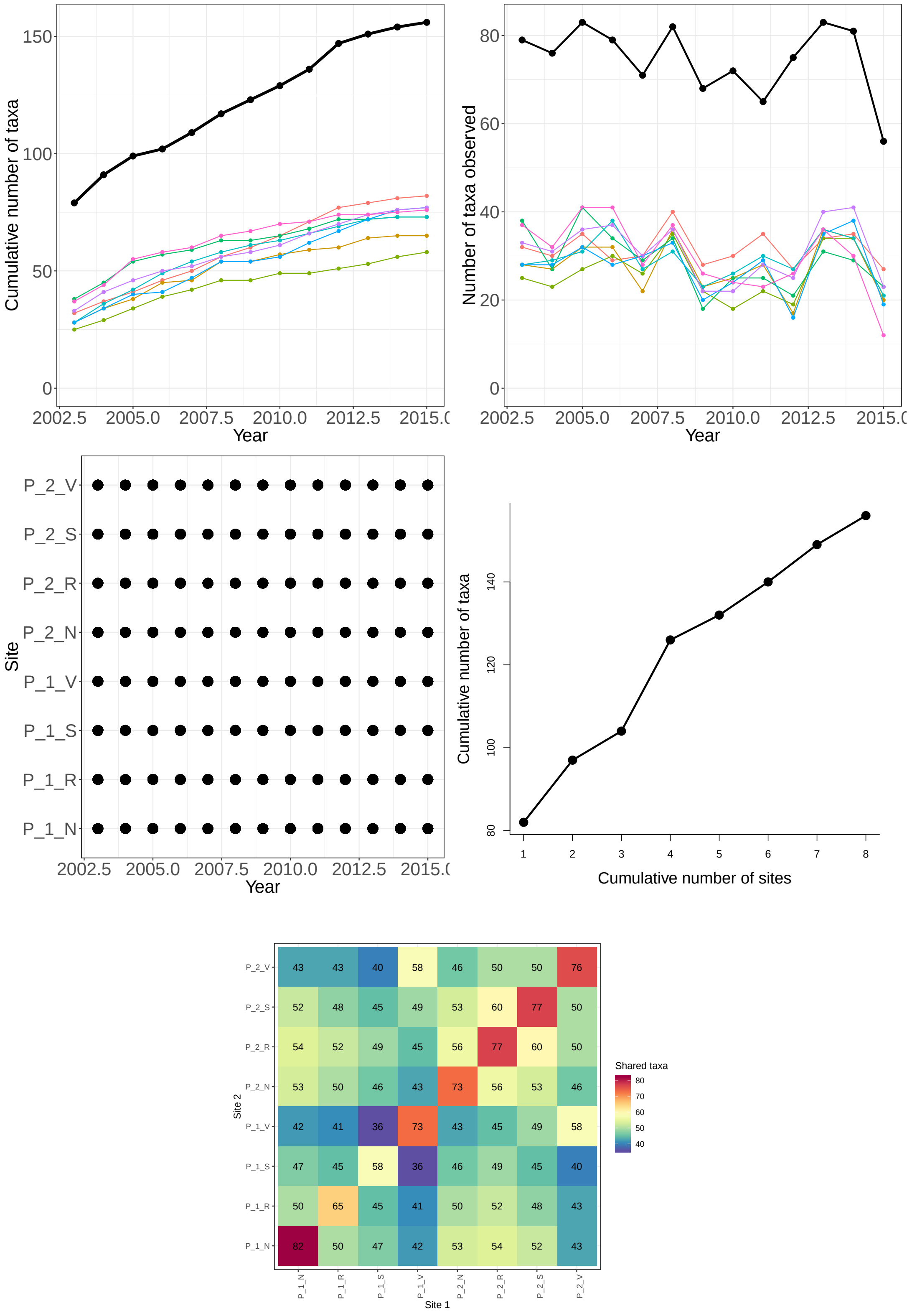


Figure 10: **SEV-plants:** Species accumulation curves (top left), annual richness (top right), and sampling effort (bottom) for plant species observed in the Pinon-Juniper habitat at the Sevilleta (SEV) LTER. The black lines represent total site-level values across all plots.

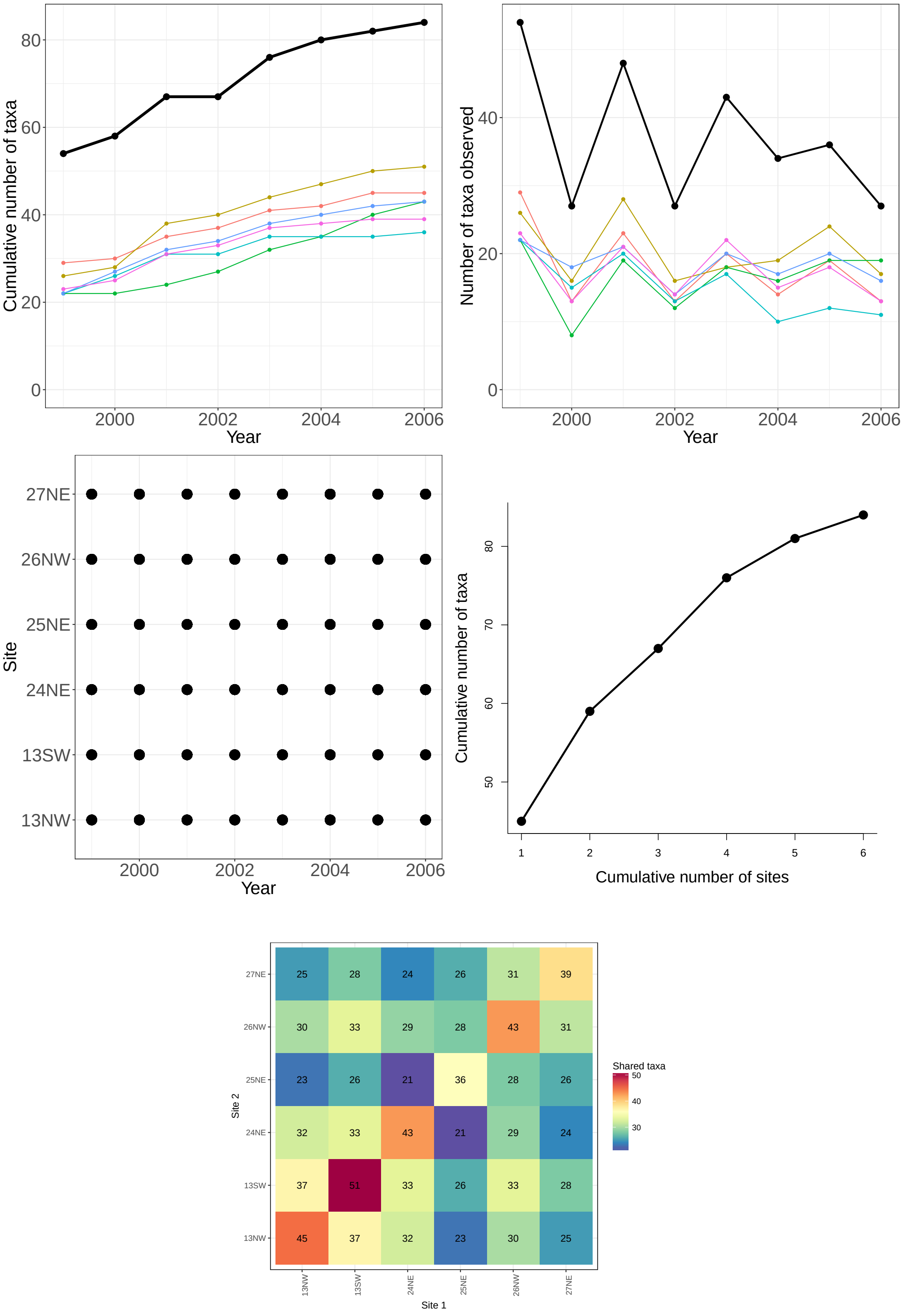


Figure 11: **SGS-plants:** Species accumulation curves (top left), annual richness (top right), and sampling effort (bottom) for plant species observed at the Shortgrass Steppe (SGS) LTER. The black lines represent total site-level values across all plots.

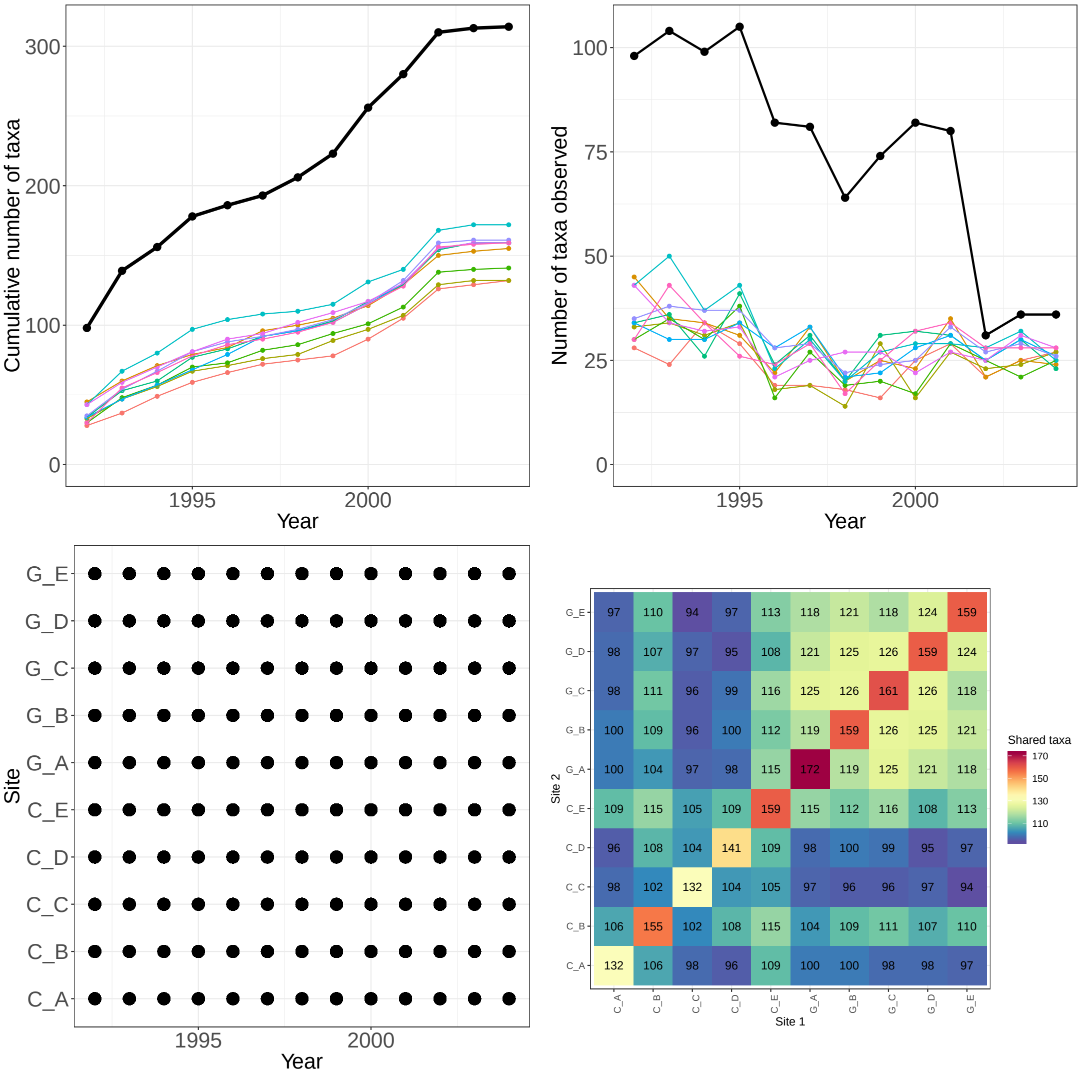


Figure 12: **SEV-arthropods:** Species accumulation curves (top left), annual richness (top right), spatio-temporal sampling effort (bottom left), and number of shared species (bottom right) for arthropod species observed in the Sevilleta LTER. The black lines represent total site-level values across all plots.

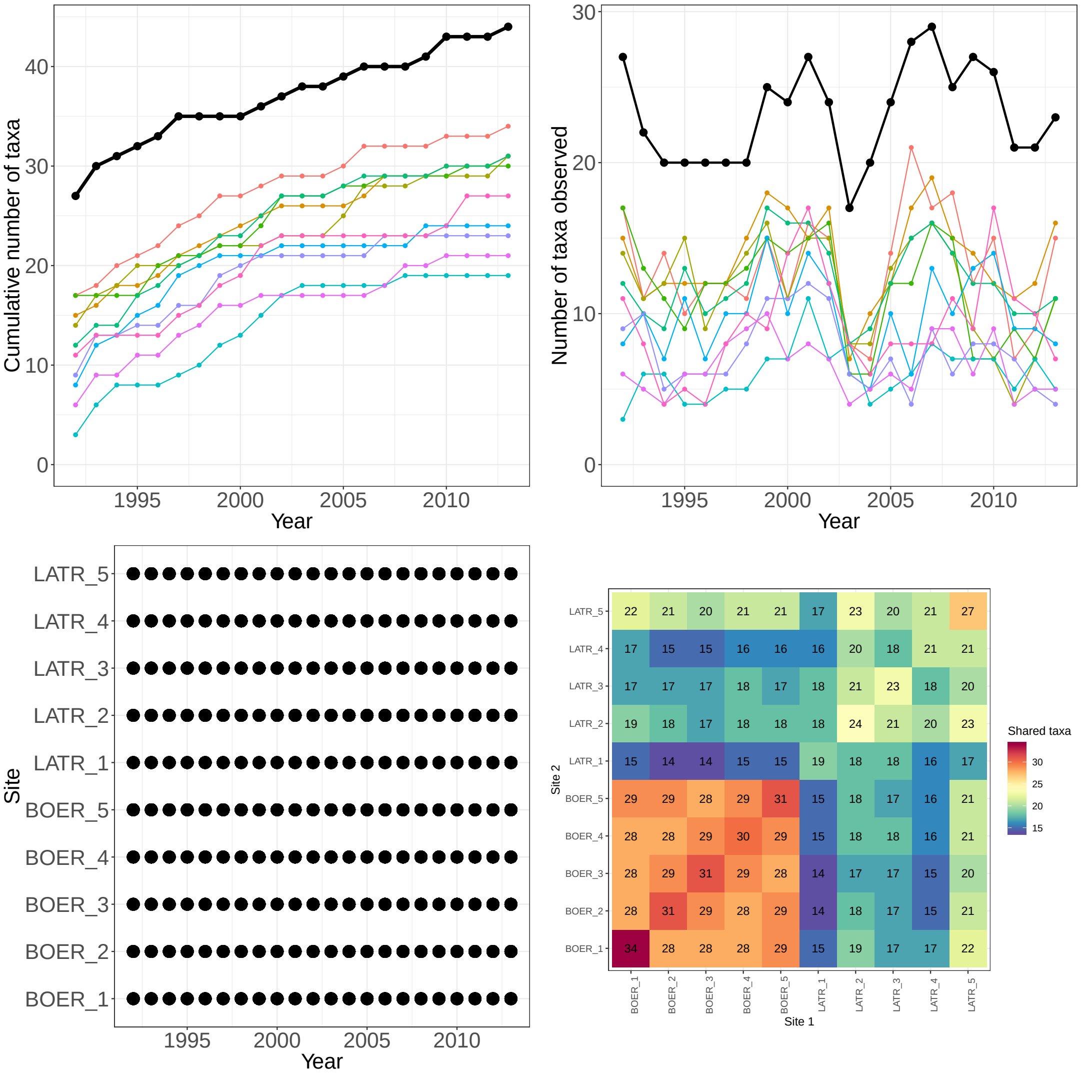


Figure 13: **SEV-grasshoppers:** Species accumulation curves (top left), annual richness (top right), spatio-temporal sampling effort (bottom left), and number of shared species (bottom right) for grasshopper species observed in Black grama (BOER) and Creosotebush (LATR) habitats the Sevilleta LTER (1992-2013). The black lines represent total site-level values across all plots.

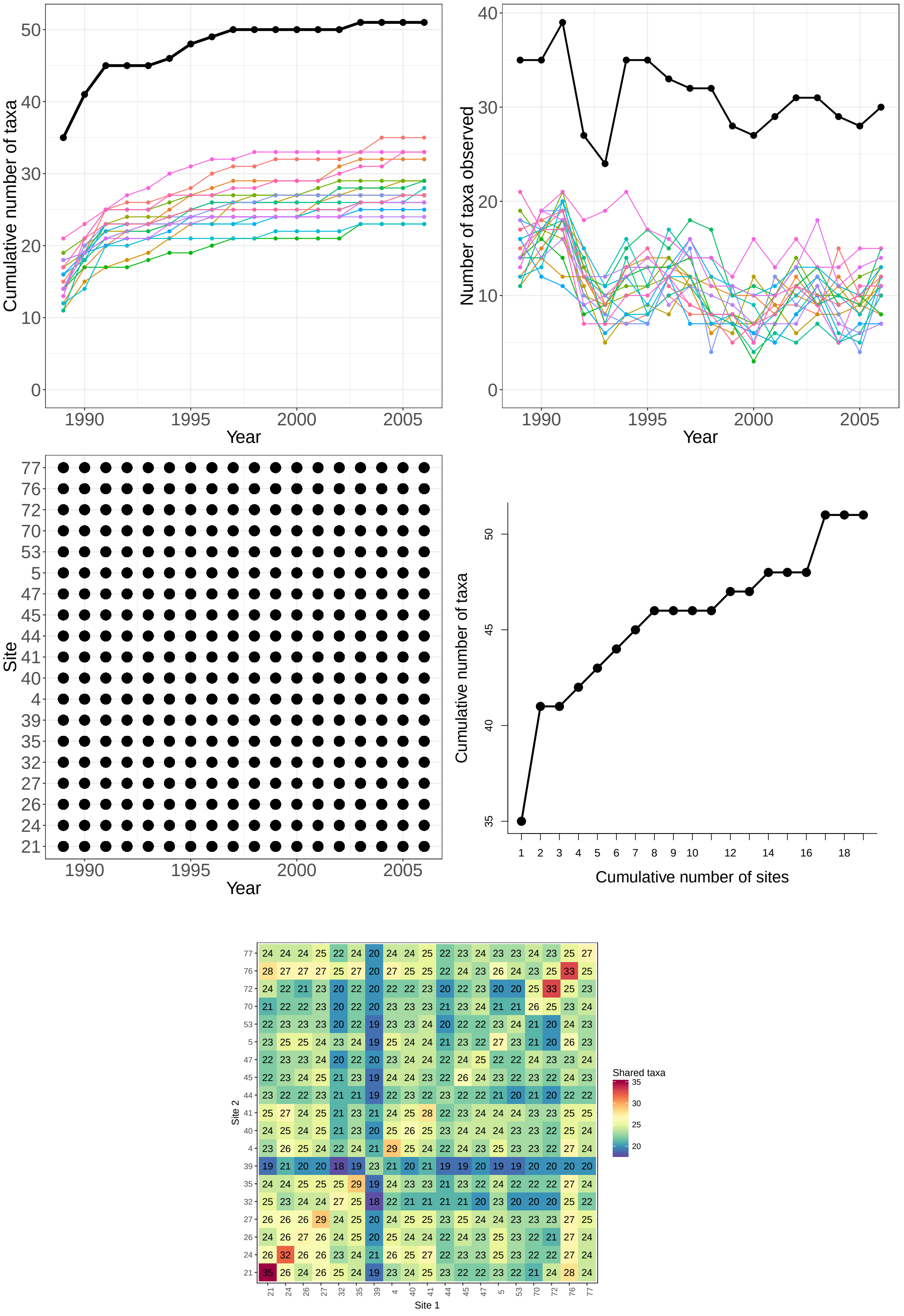


Figure 14: **CDR-grasshoppers:** Species accumulation curves (top left), annual richness (top right), spatio-temporal sampling effort (bottom left), and number of shared species (bottom right) for grasshopper species observed at the Cedar Creek LTER. The black lines represent total site-level values across all plots.

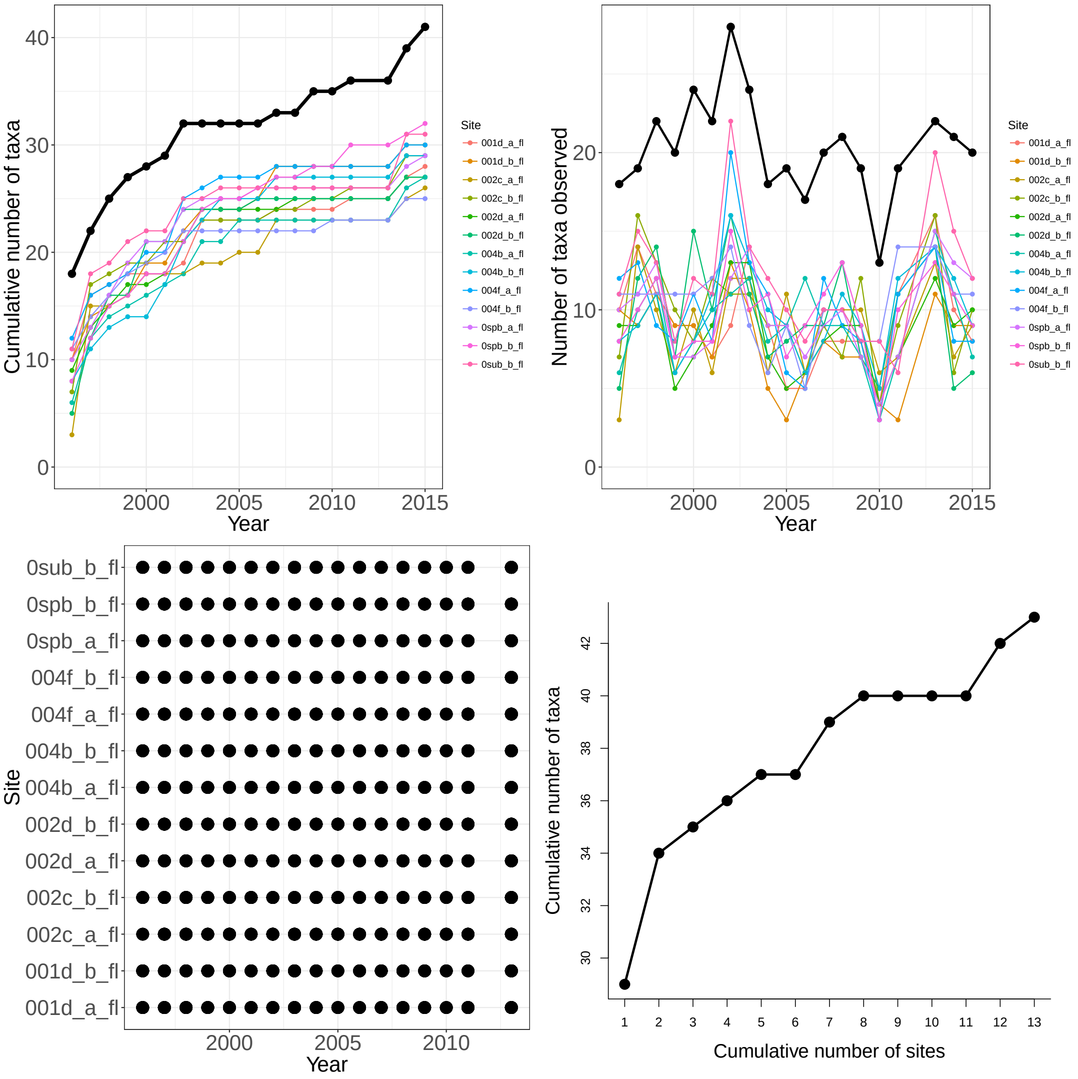


Figure 15: **KNZ-grasshoppers:** Species accumulation curves (top left), annual richness (top right), spatio-temporal sampling effort (bottom left), and number of shared species (bottom right) for grasshopper species observed in ... The black lines represent total site-level values across all plots.

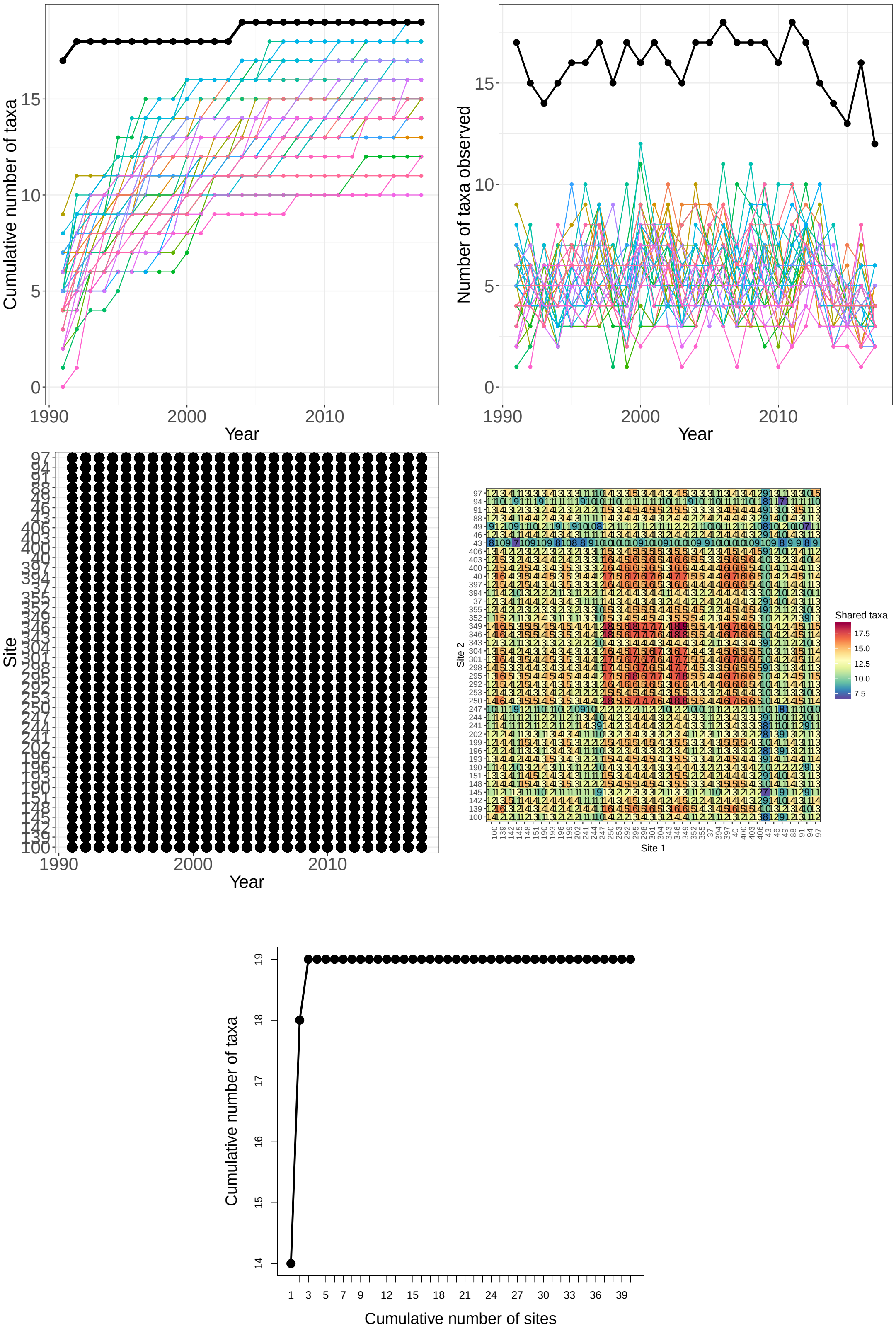


Figure 16: **LUQ-snails:** Species accumulation curves (top left), annual richness (top right), spatio-temporal sampling effort (bottom left), and number of shared species (bottom right) for snail species observed in Luquillo LTER. The black lines represent total site-level values across all plots.

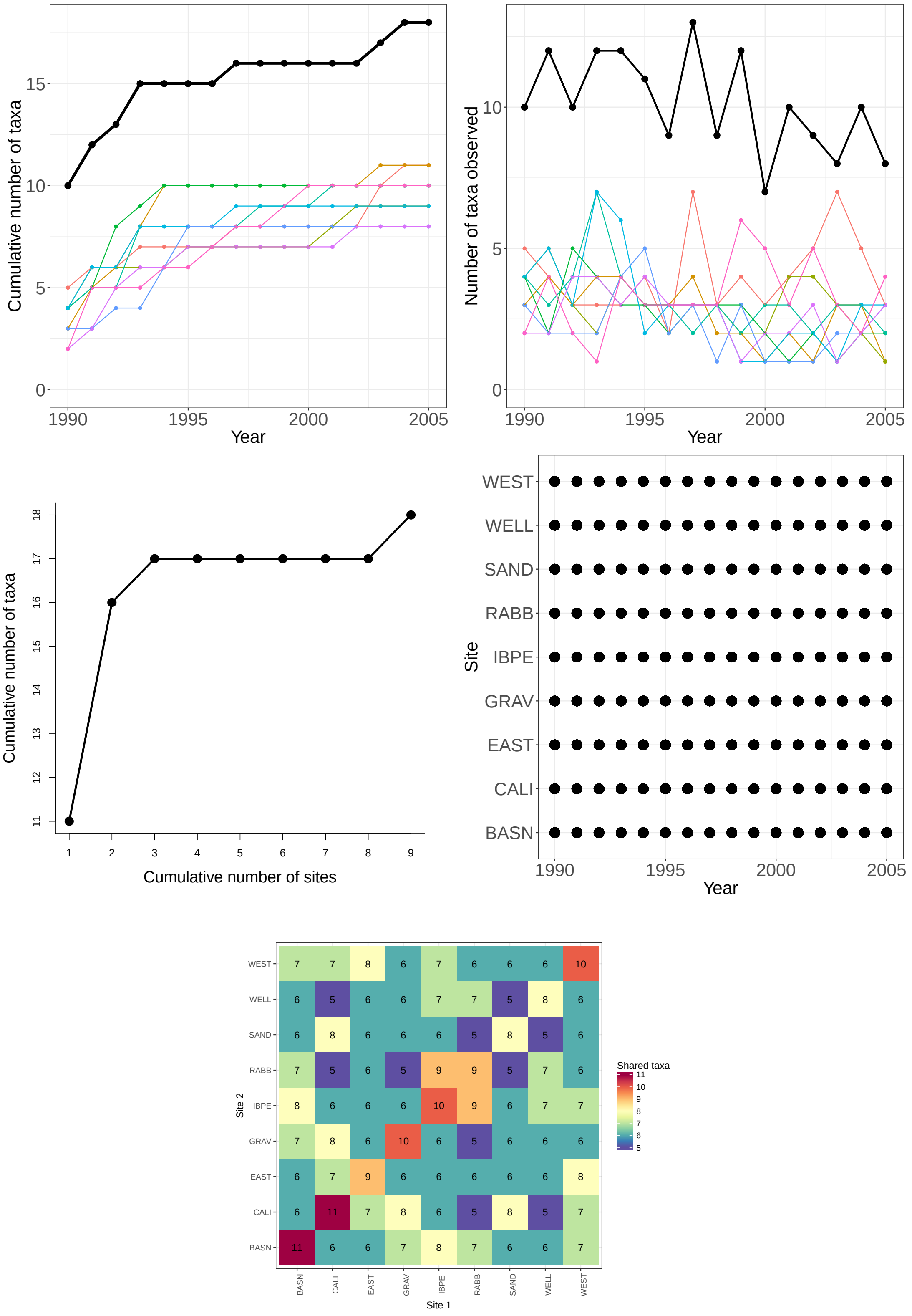


Figure 17: **JRN-lizards:** Species accumulation curves (top left), annual richness (top right), and sampling effort (bottom) for 20 lizard species observed at 9 plots in the Jornada LTER (1990-2005). The black lines represent total site-level values across all plots.

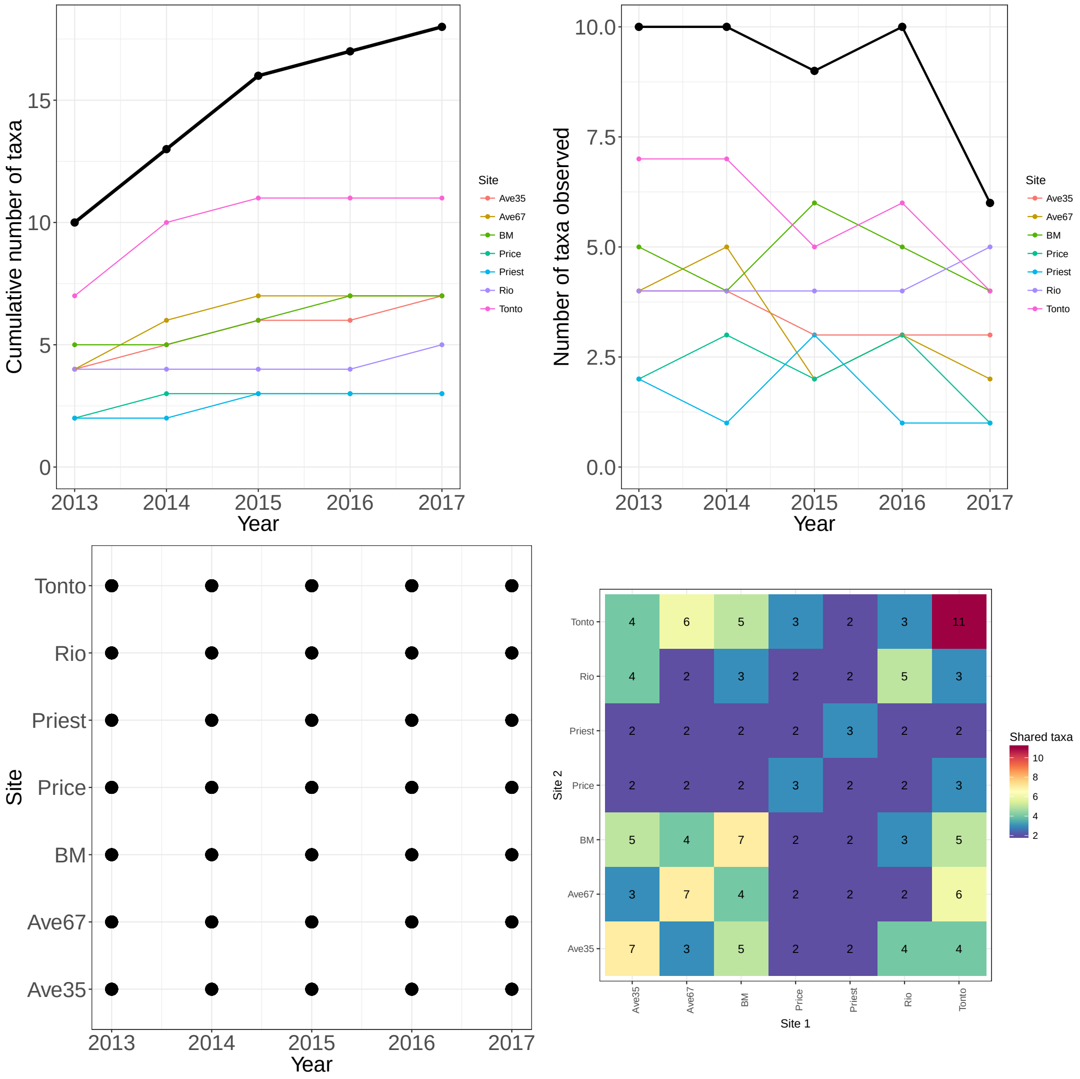


Figure 18: **CAP-herps:** Species accumulation curves (top left), annual richness (top right), and sampling effort (bottom) for species observed in the Central Area Phoenix LTER (1990-2005). The black lines represent total site-level values across all plots.

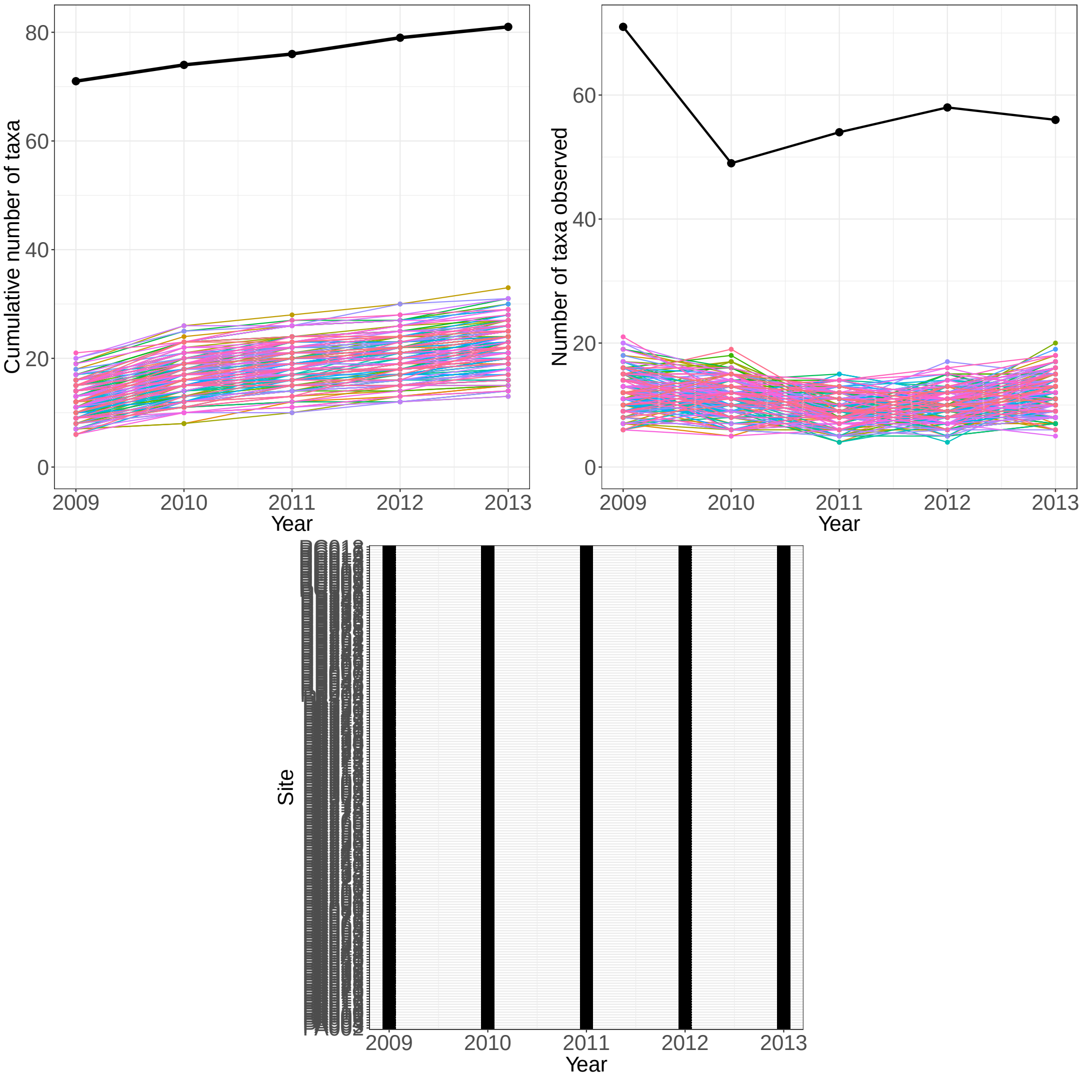


Figure 19: **AND-birds:**

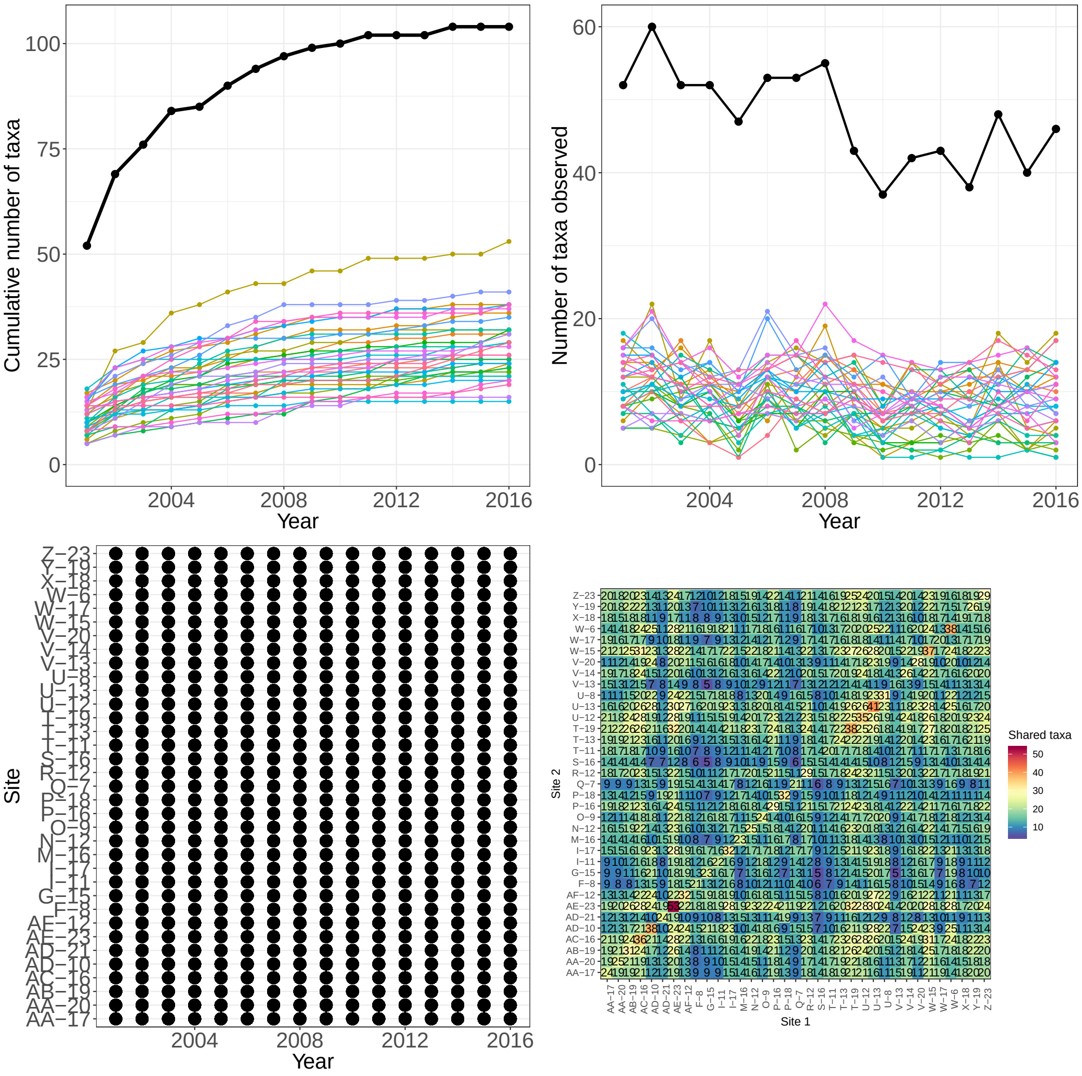


Figure 20: **CAP-birds:**

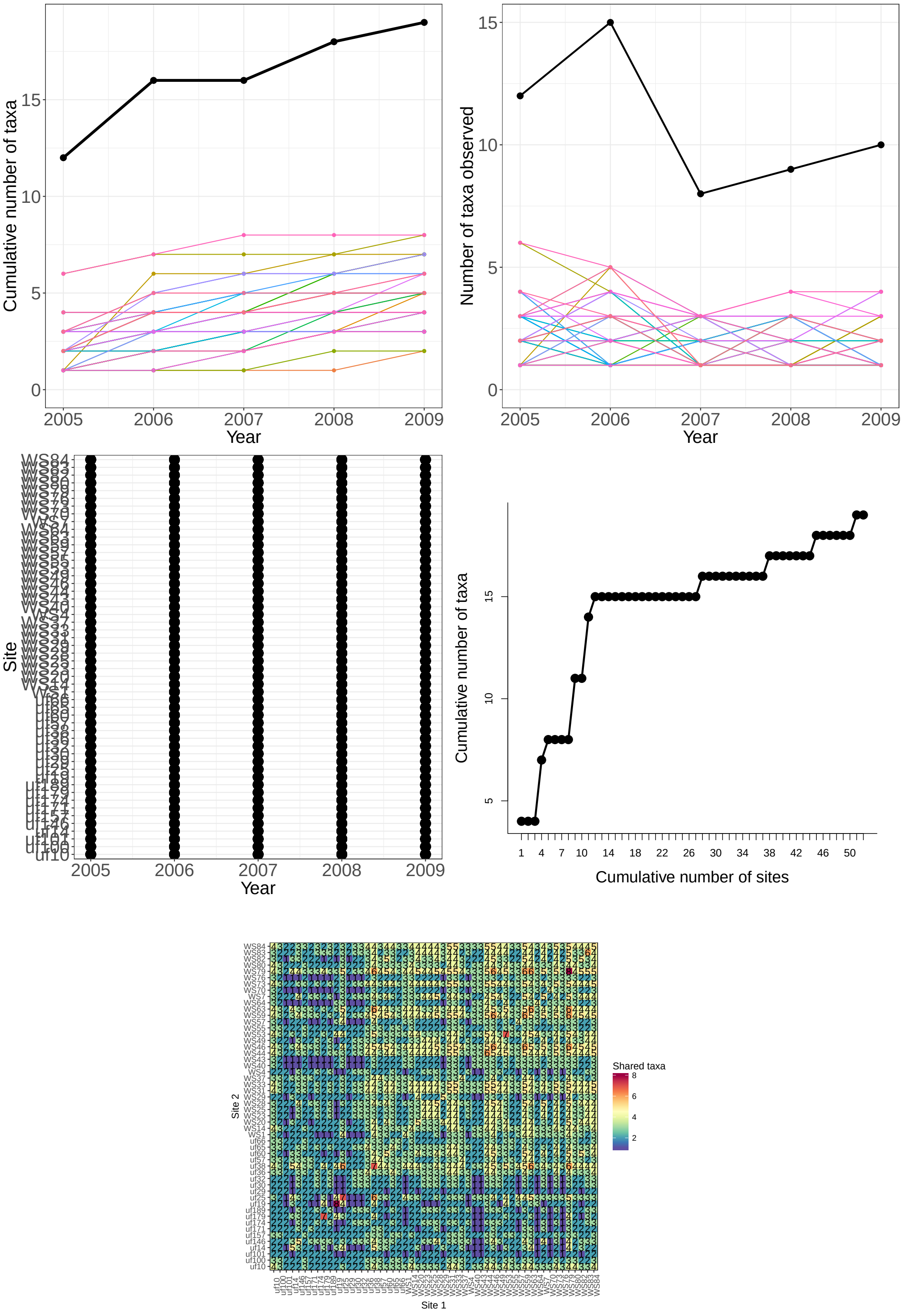


Figure 21: **BES-birds:**
